# Squamation and scale morphology at the root of jawed vertebrates

**DOI:** 10.1101/2022.02.15.480555

**Authors:** Yajing Wang, Min Zhu

## Abstract

Placoderms, the most basal jawed vertebrates, are crucial to understanding how the characters of crown gnathostomes comprising Chondrichthyes and Osteichthyes evolved from their stem relatives. Despite the growing knowledge on the anatomy and diversity of placoderms over the past decade, the dermal scales of placoderms are predominantly known from isolated material, either morphologically or histologically, resulting in their squamation being poorly understood. Here we provide a comprehensive description of the squamation and scale morphology of a primitive taxon of Antiarcha (a clade at the root of jawed vertebrates), *Parayunnanolepis xitunensis*, based on the virtual restoration of an articulated specimen by using X-ray computed tomography. Thirteen morphotypes of scales are classified to exhibit how the morphology changes with their position on the body in primitive antiarchs, based on which nine areas of the post-thoracic body are distinguished to show their scale variations in the dorsal, flank, ventral, and caudal lobe regions. In this study, the histological structure of yunnanolepidoid scales was described for the first time based on disarticulated scales from the type locality and horizon of *P. xitunensis*, demonstrating that yunnanolepidoid scales are remarkably different from the dermal plates of antiarchs including yunnanolepidoids in the absence of developed middle layer. Together, our study reveals that the high regionalization of squamation and the bipartite histological structure of scale might be plesiomorphic for antiarchs, and jawed vertebrates in general.

## Introduction

Jawed vertebrates possess a fully mineralized dermal skeleton that falls into two categories: macromery (large bony plates) or micromery (small scales), with the latter differentiating as a homogeneous squamation across the animal (Jerve et al., 2017). The squamation is a highly complex integument that varies both interspecifically and intraspecifically (Reif, 1985), and corresponds perfectly with ecology or lifestyle as well as the body plan (Ferron & Botella, 2017). It also contributes to morphological data that have been used to resolve the interrelationships of early jawed vertebrates (Choo et al., 2017; Qiao et al., 2016).

Placoderms constitute the upper part of the stem gnathostomes (Dupret et al., 2014; Qiao et al., 2016; Zhu et al., 2009). They primitively carry extensive dermoskeleton (Giles et al., 2013), but appear to have recurrently reduced or lost the dermal scales in later species including many advanced arthrodires, euantiarchs, and ptyctodonts (Denison, 1978; Janvier, 1996). Since the squamation in placoderms has only been known from a few examples (*Goujet, 1973; Gross, 1963; Hemmings, 1978; Ivanov et al., 1996; Long & Werdelin, 1986; Lyarskaya, 1977; Stensiö, 1969; Upeniece & Upenieks, 1992*), the large number of isolated placoderm scales are of uncertain affinity and little is known about the complexity of the placoderm squamation.

Concerning the histology, placoderms have been extensively studied on gnathal plates and different categories of the dermoskeleton including scales, dermal bony plates, and fin spines (Burrow & Turner, 1999; Downs & Donoghue, 2009; Giles et al., 2013; Jerve et al., 2017; Rucklin & Donoghue, 2015; Young, 2003). However, the histology of yunnanolepidoids, the most primitive group of antiarchs at the root of jawed vertebrates (Brazeau et al., 2020; Cui et al., 2019; Long et al., 2015; Zhu et al., 2016; Zhu et al., 2013), is poorly known, especially for the scales.

*Parayunnanolepis xitunensis*, a yunnanolepidoid antiarch from the Lower Devonian of South China, is known for preserving most of the squamation articulated with the dermal armor (*Figure 1A–C*). As the only near-completely known fish among yunnanolepidoids, it was regularly involved in the phylogenetic analyses of early jawed vertebrates as a representative of primitive antiarchs (Brazeau et al., 2020; Chevrinais et al., 2017; Dearden et al., 2019; Giles et al., 2015; Ribeiro et al., 2021; Zhu et al., 2021). The scale morphology of *P. xitunensis* was briefly described (Zhang et al., 2001; Zhu et al., 2012). Nevertheless, the morphological disparity of scales, which would enhance our understanding of the ancestral conditions of jawed vertebrates, was almost unremarked. In the present study, we classify and describe the scales of *P. xitunensis* and establish a squamation model for primitive antiarchs. The histology of yunnanolepidoid scales is investigated to show its deviations from that of yunnanolepidoid dermal bony plates as well as from that of euantiarch scales. To the end, we compare the histological structure and the sculpture pattern of scales across diverse groups to explore the evolution of dermal scales in early jawed vertebrates.

**Figure 1.**
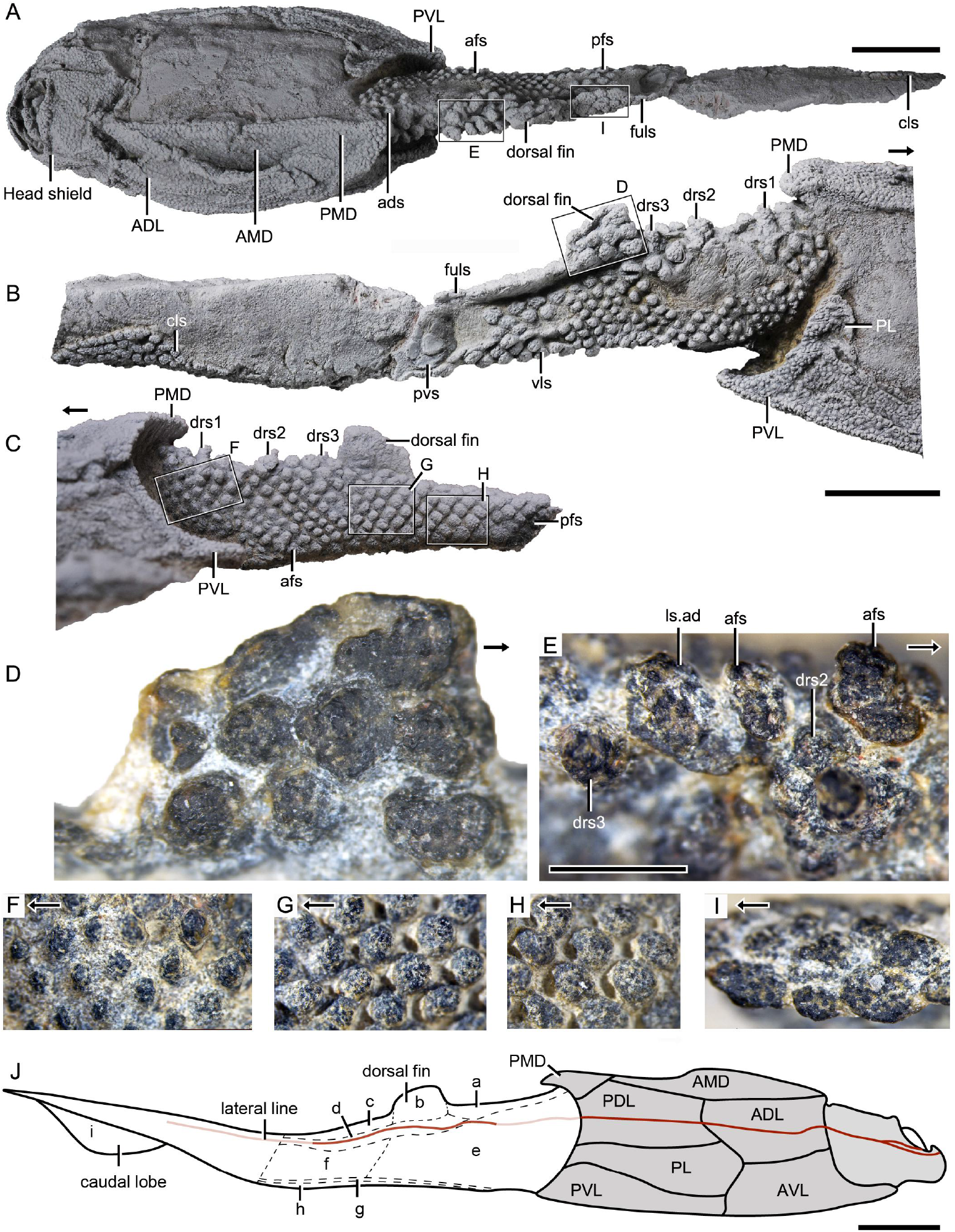
Photograph of *Parayunnanolepis xitunensis*, holotype IVPP V11679.1. (**A**) Dorsal view. (**B**) Right lateral view. (**C**) Left lateral view. (**D**–**I**) Details of (**A**–**C**) are indicated by rectangles. (**J**) Squamation model of *P. xitunensis*. Small letters in (**J**) represent the (a) predorsal, (b) dorsal fin, (c) posterodorsal, (d) dorsolateral, (e) anterior flank, (f) posterior flank, (g) ventral, (h) ventrolateral and (i) caudal lobe areas. The black arrow indicates the anterior direction. Deep and light red lines represent the exact and inferred trajectory of the lateral line, respectively. Scale bars in **A**–**C** and J equal to 5 mm, in **D**–**I** equal to 1 mm.

## Results

### Scale morphology

The post-thoracic body is completely covered by scales with substantial variability in shape (oval, or rhombic, or polygonal), size (length ranging 0.3–2.6 mm, width ranging 0.2–2.3 mm, height ranging 0.1–0.4 mm), overlap relationship (largely overlapping or no overlap), crown/base proportion, and base morphology (bulging, or flat, or concave). The vast majority of the scales are rhombic in shape with their base concave and larger than the crown. Discrete tubercles ornament the scale crown and the tubercle size changes with the different scale varieties (*Figure 1D–I*). The available scales were provisionally assigned to the thirteen morphotypes here.

*Morphotype 1*— anterior dorsal scale (*Figure 2H—figure supplement 1A–E*). The scale is distinguishable by a thickened crown, rounded anterior margin, and broad outline (width/length, abbreviated as w/l, ranging 1.3–1.6). The depressed field, i.e. the area underlying neighboring scales (Chen et al., 2012), occupies 28%–43% of the scale length. The base is concave under the crown, but convex under the depressed field. The left scale overlaps the right and the back ones (*Figure 2I, J*). The shape changes from roughly pentagonal at the first row to triangular at the second row, along with the size getting larger. Following the functional summary for various scale types by Reif (1985), the thickened crown might protect the anterior dorsal scale from abrasion in contact with the posterior median dorsal plate (PMD) of the trunk shield.

**Figure 2.**
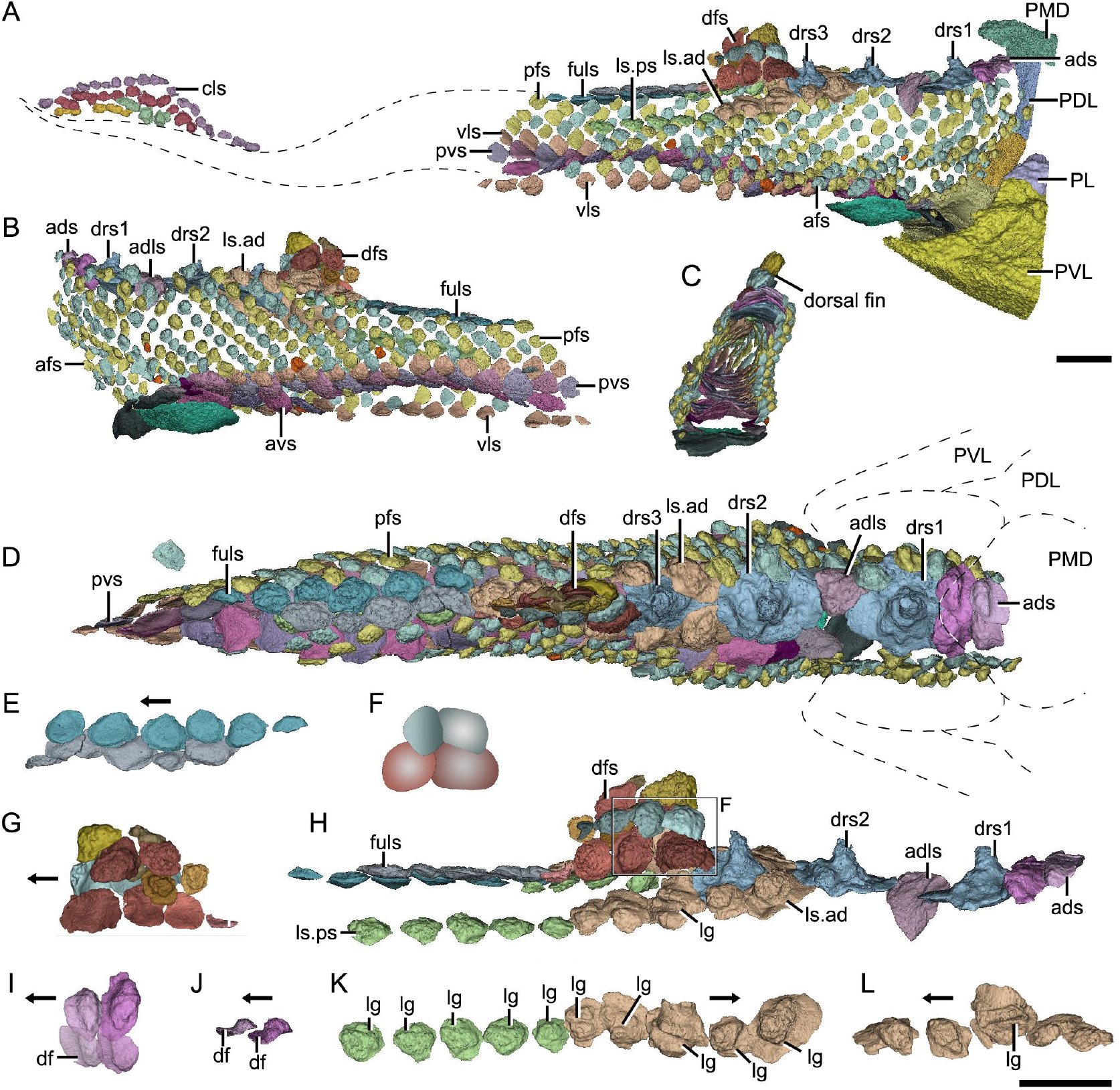
with 2 supplements. Reconstructed squamation of *P. xitunensis* based on CT scanning, IVPP V11679.1. (**A**–**D**) Squamation in (**A**) right lateral, (**B**) left lateral, (**C**) anterior, and (**D**) dorsal views. (**E**) Posterodorsal fulcral scales in ventral side. (**F**) Interpretative diagram of the dorsal fin scales, showing their contact relationships. (**G**) Dorsal fin scales in left lateral view. (**H**) Dorsal scales in right lateral view. (**I**–**J**) Anterior dorsal scales in (**I**) dorsal and (**J**) left lateral views. (**K**) Morphotype 6–7 scales on the right side, showing the trajectory of the lateral line groove. (**L**) Morphotype 6 scales on the left side. The black arrow indicates the anterior direction. Scale bars equal to 2 mm.

*Morphotype 2*—dorsal ridge scale (*Figure 2D—figure supplement 1F, H–I*). Also termed the ‘dorsal scute’ or ‘median dorsal scale’ (Zhang, 1978; Zhu et al., 2012), it is represented by three large, symmetric, polygonal scales (w/l of about 0.8) behind the anterior dorsal scales. Each scale bears a conspicuous bump. The width of the scale decreases backward, with the bump sitting more posteriorly relative to the center of the scale from the first to the third one. The first dorsal ridge scale is featured by a roughly quadrangular outline and a vaulted base. The second one has waved lateral margins and a nearly flat base. The third one, narrow heptagonal in shape, bears developed anterior and lateral corners, and a slender longitudinally directed ridge (*Figure 2H—figure supplement 1I*, dr). The base of the third scale is deeply concave beneath the ridge.

*Morphotype 3*—anterior dorsolateral scale (*Figure 2H—figure supplement 1G*). Lying between the first and second dorsal ridge scales, the scale has a quadrilateral outline (w/l of 1.17) with a protruding lateral corner. A thickening at the turn of the dorsal and ventral laminae strengthens the scale. A depressed field, which is covered by anterior flank scales, occupies almost a half-length of the scale.

*Morphotype 4*—dorsal fin scale (*Figure 2H—figure supplement 2*). The scales are probably arranged into at least three horizontal files on either side of the dorsal fin, with up to four scales on each file (Figures 1D and 2G). They change the outline and decrease progressively in size towards both the dorsal and trailing edges of the fin. In the basal horizontal file, the back scale overlaps the front one, while in the second file, the front scale overlaps the back one (*Figure 2F*), revealing the variability of scale overlap relationships.

*Morphotype 5*—posterodorsal fulcral scale (*Figure 2D—figure supplement 1Q–T*). The scale is rostrocaudally elongated, about 1.2–1.8 times longer than width, and sub-rectangular in shape. The scales in two files are arranged in a staggered manner (Figure 1I), with the left underlying the right. Remarkably, the base of the scale in the left file is concave, while that in the right is convex (*Figure 2E*).

*Morphotype 6*—anterodorsal lateral line scale (*Figure 2H—figure supplement 1J–N*). The scales are arranged in an arcuate file around the base of the dorsal fin. They vary in morphology along the body: the first three scales are featured by a high crown, while the last two by a relatively flat crown; their shape changes from oval to rhombic (w/l of 0.7–1.0). The lateral line canal passes around the crown periphery of the first two scales, but through the crown surface of the following three ones as an open groove (*Figure 2K, L, lg*).

*Morphotype 7*—posterodorsal lateral line scale (*Figure 2H—figure supplement 1O, P*). The scale is rhombic in shape with a size intermediate between adjacent flank and fulcral scales. The base is concave and shows a thickening centrally. The scales also carry a lateral line groove that extends from the Morphotype 6 scales and parallels to the longitudinal axis of the body.

*Morphotype 8*—anterior flank scale (*Figure 3A—figure supplement 3A–E, J–N*). The scales are oval to sub-rhomboid in shape and form the first 15 (on the right side) or 18 (on the left side) rows (*Figure 3A, B*) on the post-thoracic body. The crown, which is narrower than the base, slightly leans back, and usually beyond the base. The first three rows consist of closely packed scales, while the following rows consist of loosely packed scales. There is a trend of scale size reduction in the top-down direction. A shallow groove on the crowns of three adjacent flank scales in rows 7–9 of the left side represents the anterior elongation of the lateral line on the Morphotype 6 scales (*Figure 3A—figure supplement 3B, C*).

**Figure 3.**
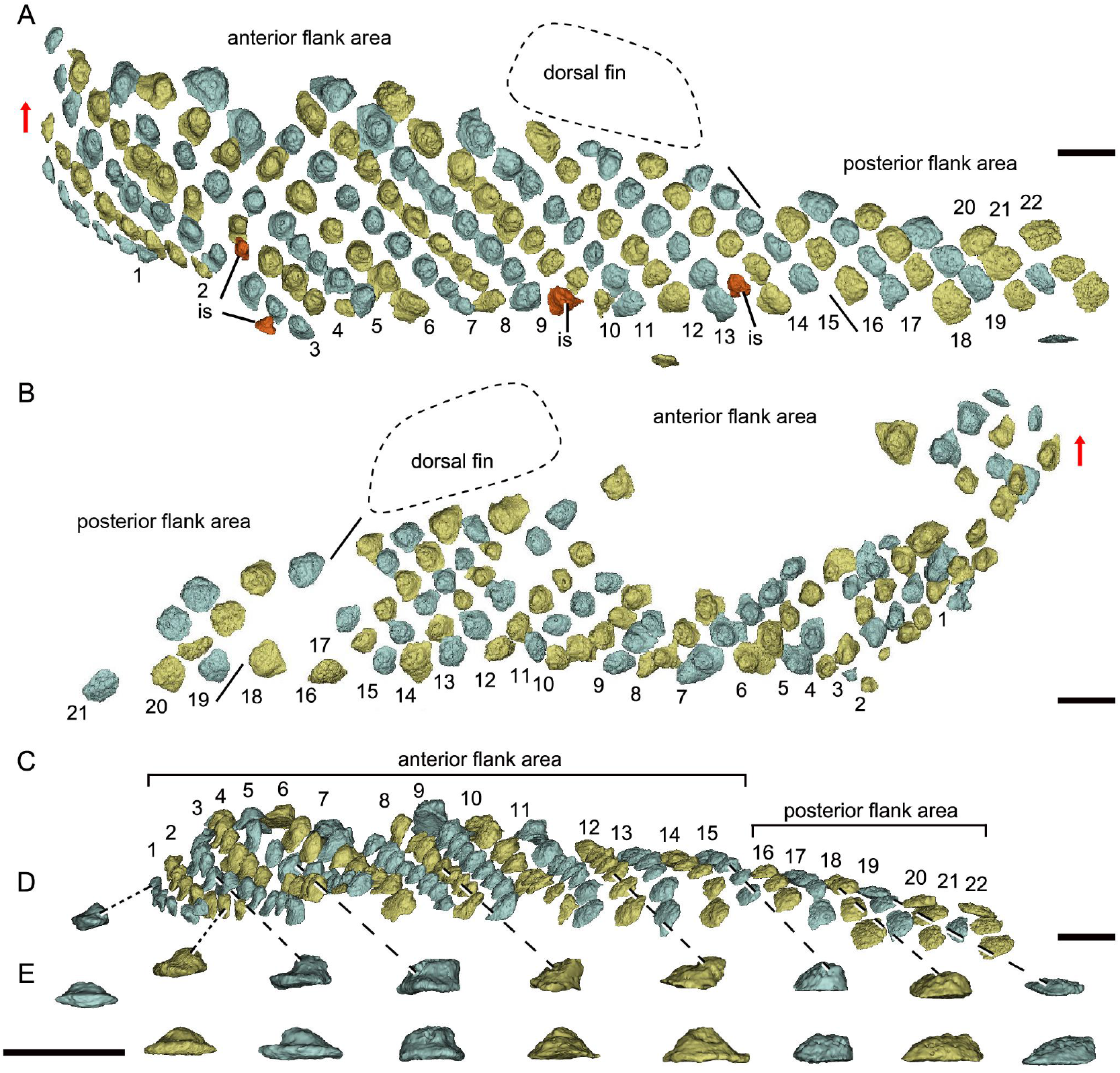
with 1 supplement. Reconstructed flank squamation of *Parayunnanolepis xitunensis* based on CT scanning, holotype IVPP V11679.1. (**A**) Left flank scales in lateral view. (**B**) Right flank scales in lateral view. (**C**) Left flank scales in ventral view. (**D**–**E**) A series of flank scales in (**D**) ventral and (**E**) anterior views, showing the anterior–posterior gradation of scales. The red arrow indicates the dorsal direction. Scale bars equal to 1 mm.

*Morphotype 9*—posterior flank scale (*Figures 3A—figure supplement 3F, I*). They are represented by the flank scales lying in rows 16–22 (on the left side) or 18–22 (on the right side), posterior to the trailing edge of the dorsal fin (*Figure 3A, B*). The scales are rectangular (w/l of 0.83) in shape. The flat crown almost shelters the base, rarely leaving a depressed field.

*Morphotype 10*—anterior ventral scute (*Figure 4A—figure supplement 4A–G*). The ventral scutes in the first to third rows are asymmetric in shape with a straight mesial margin. The crown of the first scute is separated from the base posteriorly by a constricted groove (*Figure 4E–G*, gr). Each scute consists of the lateral and ventral laminae. The right scute overlaps the left one in the first two rows, but the left one in the third row overlaps the right one (*Figure 4D, H, I*).

**Figure 4.**
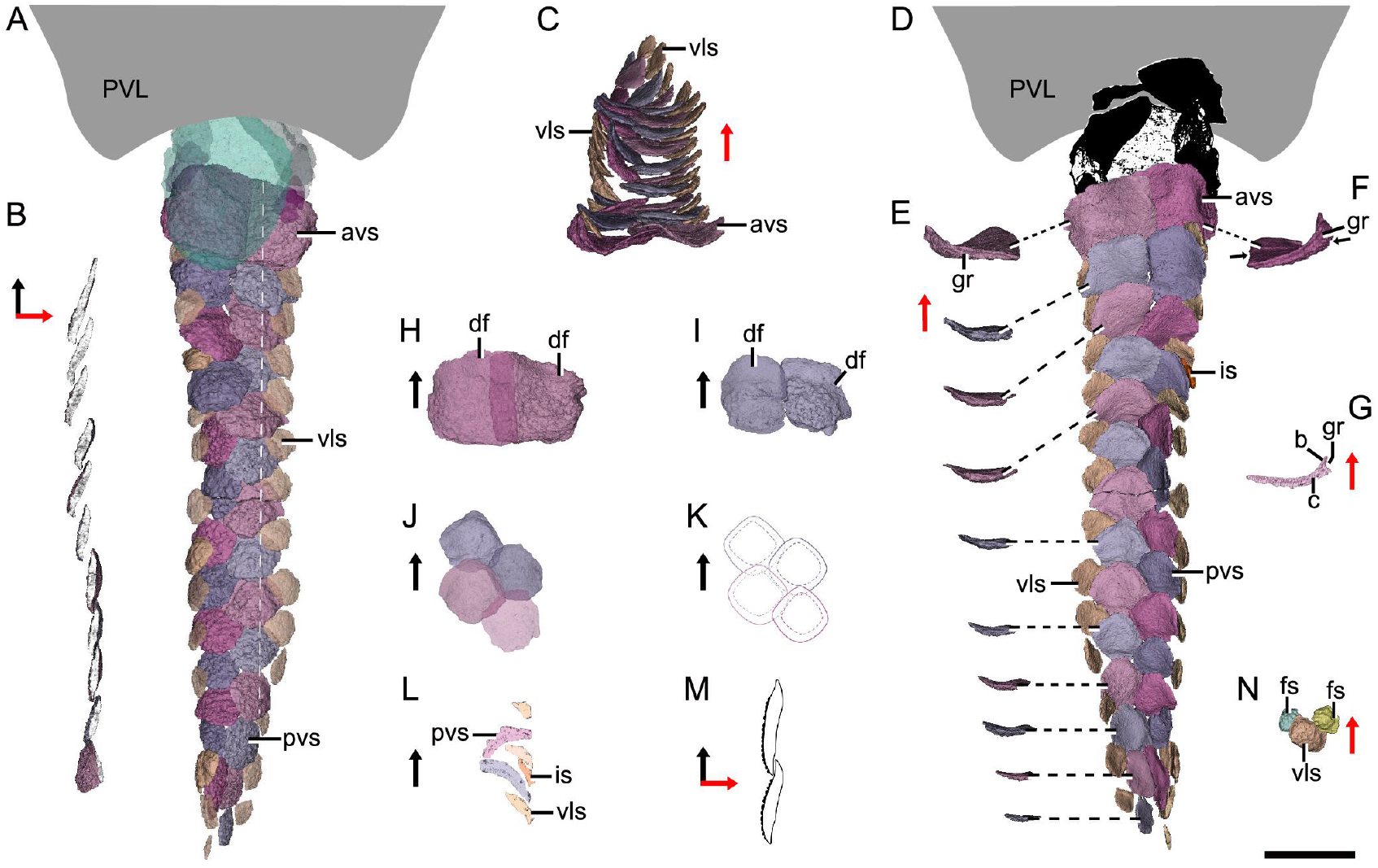
with 1 supplement. Reconstructed ventral squamation of *Parayunnanolepis xitunensis* based on CT scanning, holotype IVPP V11679.1. (**A**) Ventral scales in ventral view. (**B**) Virtual axial section of the ventral scutes at the level indicated by the white dotted line in (**A**). (**C**) Ventral scutes in anterior view. (**D**) Ventral scales in dorsal view. (**E**) A series of ventral scutes in posterior view. (**F**) The first right ventral scute in posterior view. (**G**) Virtual axial section of the scale in (**F**). (**H**–**I**) Anterior ventral scutes in (H) first and (I) second rows in ventral views. (**J**–**K**) Reconstruction and interpretative reconstructions of the posterior ventral scutes in the fourth and fifth rows in ventral view, respectively. (**L**) Virtual sagittal section of the additional scale and its surrounding scales. (**M**) Schematic diagram of the articulated way between ventral scutes in the front and back rows. (**N**) Ventrolateral scale and its surrounding flank scales. The black and red arrows indicate the anterior and dorsal directions, respectively. Scale bar equal to 2 mm.

*Morphotype 11*— posterior ventral scute (*Figures 4D—figure supplement 4J–N*). The ventral scutes behind the third row share a staggered arrangement (*Figure 4J, K*). Each scute overlaps the two back ones. The scute is nearly symmetric, with a roughly rhombic shape in most cases. The base is thickened and concave centrally. The upturned posterior margin of the front scute closely hooks the downturned anterior margin of the back one (*Figure 4B, M*). Compared with Morphotype 10, the lateral lamina in the Morphotype 11 scale is much less developed (*Figure 4C, E*).

*Morphotype 12*—ventrolateral scale (*Figure 4A—figure supplement 4H, I*). The scale is intermediate in size and morphology between the flank scales and ventral scutes. It is characterized by a rhombic shape (w/l of 0.9–1.1), with the mesial corner always leveling in front of the lateral corner. A narrow overlapping occurs between the Morphotype 12 scale and its adjacent ones (*Figure 4D, N*).

*Morphotype 13*—caudal lobe scale (*Figure 2A*). The scales in a small rectangular or square shape cover the hypochordal lobe. They align into four parallel rows in a longitudinal direction.

### Squamation

The post-thoracic body is roughly triangular-shaped in the transverse section (*Figure 2C*). It is broadest at the level just posterior to the ventral wall of the trunk shield (*Figure 2D*), with its foremost portion stretching into the trunk shield. The dorsal fin is positioned at about the first third of the post-thoracic body length (*Figures 1B* and *2A, B*), and has its preserved part of about 3.0 mm in length, 1.0 mm in width, and 1.6 mm in height.

The scales are packed into oblique dorsoventral rows (flank scales), longitudinal files (dorsal and ventral scales), and curved linear rows (caudal lobe scales)(*Figure 5—figure supplement 5*). According to the scale morphotypes and arrangements, the squamation can be divided into four large regions (dorsal, ventral, flank, and caudal lobe regions), which can be further divided into nine areas (*Figure 1J*).

**Figure 5.**
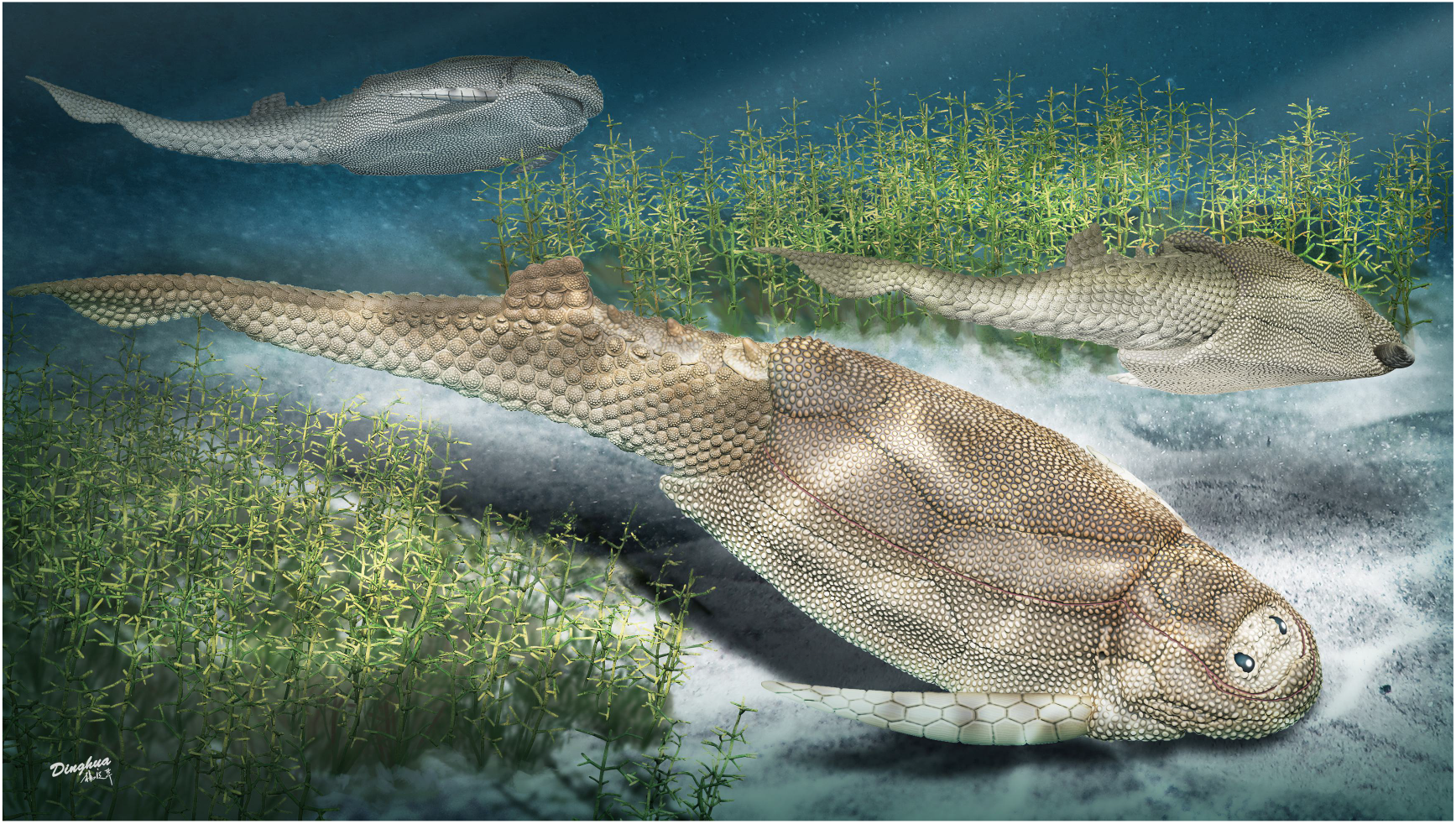
with 1 supplement. Life reconstruction of *Parayunnanolepis xitunensis*, drawn by Dinghua Yang.

The dorsal region comprises the predorsal, dorsal fin, posterodorsal, and dorsolateral areas. The squamation in the predorsal area shows the large disparity of scales (Morphotypes 1–3), which are more heavily imbricated with each other (*Figure 1E*) than the scales in the other areas. Three specialized dorsal ridge scales (Morphotype 2) lie in sequence just behind the anterior dorsal scales (Morphotype 1) but in front of the dorsal fin. The first two dorsal ridge scales are separated by a pair of anterior dorsolateral scales (Morphotype 3). The dorsal fin area is completely covered by numerous imbricating scales (Morphotype 4)(*Figure 2G, H*), a feature also known in other placoderms with heavy scale cover. In the posterodorsal area, two files of flat fulcral scales (Morphotype 5) are positioned just behind the dorsal fin. The main lateral line is discernible as an open groove running around or through the crowns of the Morphotype 6–7 scales in the dorsolateral area.

The flank region comprises the anterior and posterior flank areas, which are covered by anterodorsally inclined (c.40–50°) rows of scales (Morphotypes 8–9), with a different row number on either side of the body (*Figure 3A, B*). The main lateral line in the dorsolateral area extends anteriorly onto the Morphotype 8 scales of the anterior flank area. A small pore opens onto the crown surface in some flank scales (*Figure 3A—figure supplement 3G, M*), which is suggestive of the mucus-producing system as presumed for that in other fishes (Turner, 1991). Remarkably, the squamation in the flank region exhibits a backward gradation of scales in shape (from oval, to rhombic or rectangular), thickness (from high to low crown), and size (from small to large in overall) (*Figures 3C–E* and *5*). The size gradation is contrary to that in many other early vertebrates, such as *Tremataspis* (Märss et al., 2015), *Guiyu* (Cui et al., 2019), *Mimipiscis* (Choo, 2012), and *Gogosardina* (Choo et al., 2009).

The ventral region comprises the ventral and ventrolateral areas. The scales in the ventral region are flatter than those in other areas. A file of ventral scutes (Morphotype 10–11) on either side of the ventral area, is flanked by a file of ventrolateral scales (Morphotype 12) in the ventrolateral area. The squamation in the ventral region also exhibits a backward gradation of scales in shape (from pentagonal to rhombic), size (from large to small), the development of lateral lamina (from well to weakly developed), and overlapping degree (from heavily to slightly overlapped) (*Figure 4A, D*). A few additional scales are interspersed between the regularly arranged ventral scutes and flank scales (*Figure 4D, L*, is; *Figure 3A—figure supplement 3K, L*). The same condition was also found in the squamation of *Asterolepis* (Ivanov et al., 1996).

The caudal lobe region, or the ventral lobe of the caudal peduncle, is covered by tiny scales (Morphotype 13). Being the smallest amongst the squamation (*Figure 5*), these scales are arranged in linear rows as the tail scales of other antiarchs, such as *Pterichthyodes* (Hemmings, 1978).

### Histology of yunnanolepidoid scales

Despite yunnanolepidoid antiarchs displaying abundant fossil records in the Early Devonian sediments of South China (Burrow et al., 2000; Zhao & Zhu, 2010; Zhu, 1996; Zhu et al., 2000), the identification of isolated scales has proven difficult as the scale sculpture, mainly composed of round tubercles, is less diagnostic. As such, yunnanolepidoid scales have never been investigated histologically. For the histological research, we selected dozens of isolated placoderm scales from the type locality and horizon of *P. xitunensis*. These isolated scales are difficult to be referred to as any definite species, however, they can be assigned to yunnanolepidoids with a reference to the articulated specimen of *P. xitunensis* described here in detail. Displaying various shapes, these scales should come from different areas of the post-thoracic body and/or different species. The thin sections through different morphotypes of yunnanolepidoid scales show that the scale uniformly comprises two main divisions: a compact superficial layer, and a thick, lamellar basal layer (*Figure 6*). Thus, the histology of yunnanolepidoid scales is remarkably different from that of their dermal bony plates (Giles et al., 2013), which exhibit a three-layered structure including a cancellous spongiosa in the middle.

**Figure 6.**
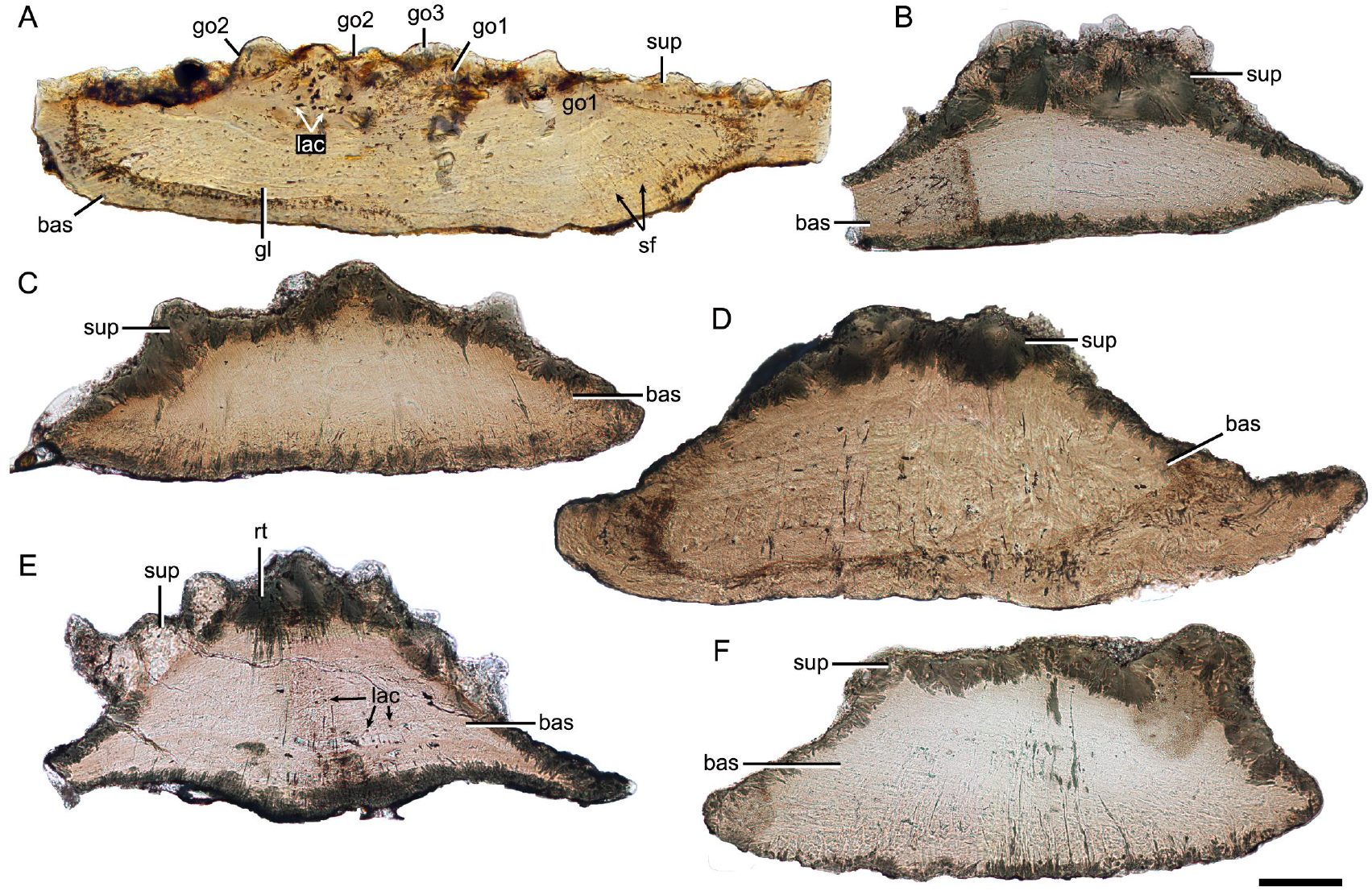
with 1 supplement. Thin sections through yunnanolepidoid scales. (**A**) IVPP V28642. (**B**) IVPP V28643. (**C**) IVPP V28644. (**D**) IVPP V28645. (**E**) IVPP V28646. (**F**) IVPP V29647. Scale bar equal to 100 μm.

The superficial layer of the yunnanolepidoid scale is confined to the tubercular ornament. Although recrystallization more or less blurs the original histological structure, the odontodes in the superficial layer display signs of superpositional stacking of at least three generations (*Figure 6A*, go1–3). The number of superimposed layers of odontodes decreases from the center of the scale to the edge. The younger odontodes cover the older generations of odontodes either completely or incompletely, suggesting a combined areo-superpositional growth (Ørvig, 1968) or a compound growth (Cui et al., 2021). Polyodontode and the compound growth pattern were ever considered to be plesiomorphic in jawed vertebrates (Cui et al., 2021; Qu et al., 2013). The conditions in yunnanolepidoid scales offer support for these hypotheses. Most of the lacunae are star-shaped with rounded or more elongated cell bodies (*Figure 6A, lac*), and have ramifying canaliculi in all directions. The basal layer is permeated by vertical Sharpey’s fibers. These fibers converge towards the center of the scale, at right angles to the lines of the laminae. The scale base shows a series of growth lines (*Figure 6A, gl*).

## Discussion

### Squamation of antiarchs

As described above, *P. xitunensis* exhibits remarkable morphological variability in the scales of nine body areas. Also present in *Romundina* and *Radotina* (Ørvig, 1975), another two primitive members of placoderms (Li et al., 2021; Vaškaninová et al., 2020), the high regionalization of squamation probably represents a primitive character for jawed vertebrates.

In comparison, the squamation regionalization in the ventral and dorsal regions of the body is reduced in euantiarchs (*Figure 7*). Most bothriolepidoids have almost naked bodies, such as *Bothriolepis canadensis*, in which only a small patch of square scales occurs along the lower margin of the tail (Stensiö, 1948). The exceptions are *B. gippslandiensis* and *B. cullodenensis* (Long, 1983; Long & Werdelin, 1986), which have a complete squamation with three regional divisions: the ventrolateral, dorsal fin, and flank areas. *Pterichthyodes milleri* appears to have the highest regional differentiation amongst asterolepidoids, with about six body areas, i.e., predorsal, dorsal fin, posterodorsal, flank, ventral, and caudal lobe areas (Hemmings, 1978). The squamation of *Asterolepis* can be divided into three areas: dorsal fin, dorsal, and flank areas, based on its scale morphotypes (Ivanov et al., 1995; Ivanov et al., 1996). The squamation of *Remigolepis* can also be divided into three areas: predorsal, posterodorsal, and flank areas (Johanson, 1997), or even fewer areas in the species from northern China (Pan et al., 1987).

**Figure 7.**
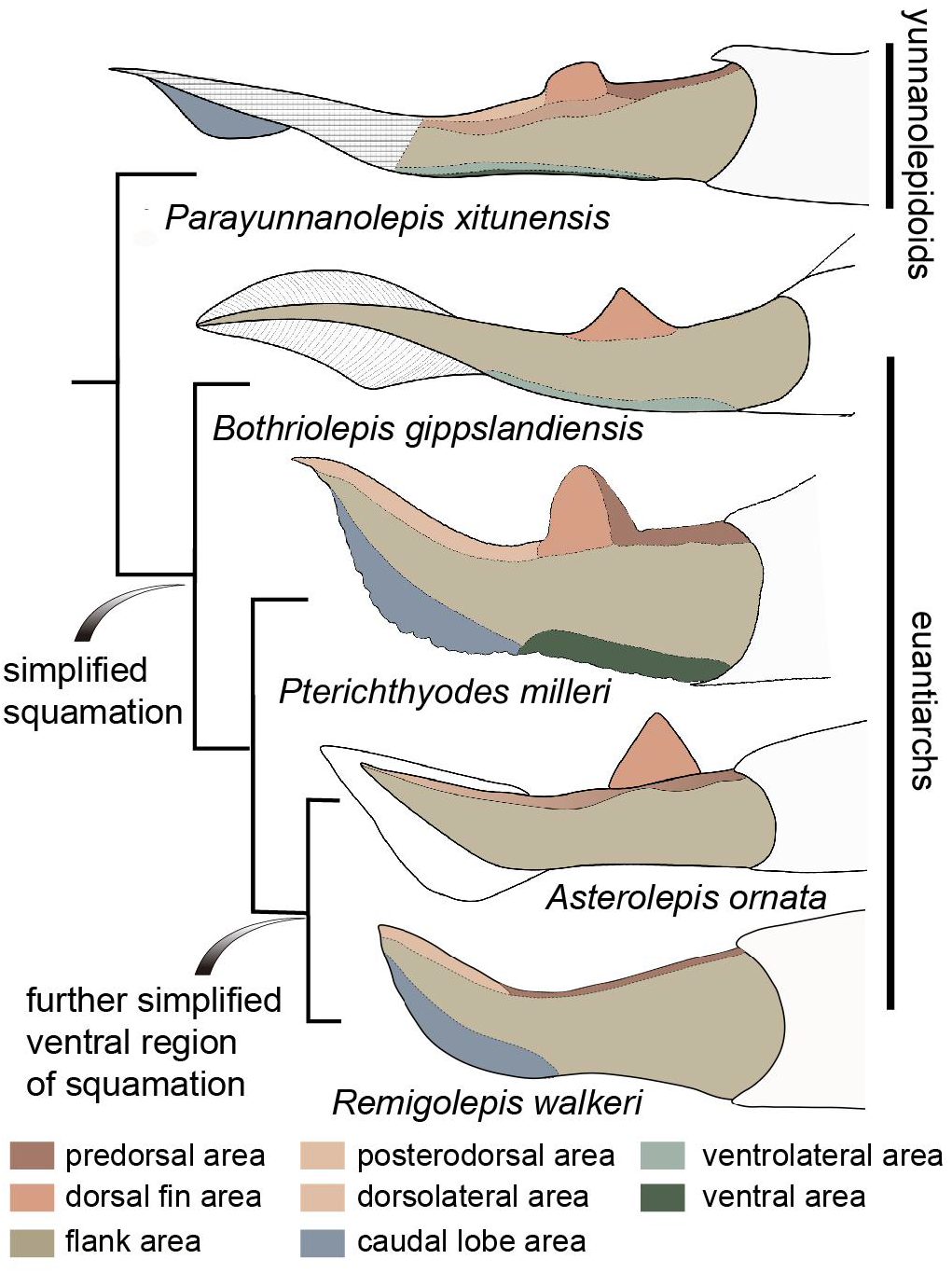
Evolution of squamation pattern in antiarchs. Redrawn from the source illustrations of Long & Werdelin (1986) for *Bothriolepis gippslandiensis*, Hemmings (1978) for *Pterichthyodes milleri*, Ivanov et al. (1996) for *Asterolepis ornata*, and Johanson (1997) for *Remigolepis walkeri*.

It seems that the squamation is simplified in the aspect of regionalization or disparity in euantiarchs, and further simplified in the clade comprising *Asterolepis* and *Remigolepis*, as their ventral scales are not differentiated from the flank scales (*Figure 7*). The simplification of squamation is also known in crown sarcopterygians and actinopterygians (Cui et al., 2019), suggesting substantial parallelism of squamation in early jawed vertebrates.

### Phylogenetic signals in scale microstructure

The dermal skeleton of heterostracans, osteostracans, placoderms, and early osteichthyans usually consists of three principal layers: a superficial layer composed of dentine and enameloid/enamel; a trabecular/cancellous layer composed of vascular bone; and a compact basal layer composed of lamellar/pseudolamellar bone (Burrow & Turner, 1999; Donoghue & Sansom, 2002; Giles et al., 2013; Qu, Haitina, et al., 2015). This tripartite structure was ever considered to be plesiomorphic for early jawed vertebrates (Qu, Haitina, et al., 2015). However, different dermal units evolve as independent modules (Qu, Haitina, et al., 2015), and vary in histological structure as suggested by the conditions in yunnanolepidoids. In this part, we compare scales of early jawed vertebrates to assess how the variation of diagnostic scale traits including histology and sculpture is correlated with the phylogenetic relationships.

In osteostracans or the immediate outgroup of jawed vertebrates, the middle layer was well described in tremataspid scales (Qu, Blom, et al., 2015). Ornament on the external surface of the osteostracan scales changes from smooth with pores, to variable shaped tubercles in combination with nodules, ridges, ribs, and inter-ridge grooves (Märss et al., 2015).

Yunnanolepidoid scales bear the round tubercle sculpture on the surface and lack the well-developed middle layer, in contrast with that of euantiarchs. The middle partition of scales assigned to *Asterolepis ornate* (Lyarskaya, 1977) and *Wurungulepis denisoni* (Burrow & Turner, 1998) coincides with the features of the middle layer, such as the “vascular canals surrounded by osteons”. The *Remigolepis* scale also develops a thick middle layer bearing the upper part of the canal system (Lukševičs, 1991). Considering ornamentation, the asterolepidoid scale is much more variable than that of yunnanolepidoid antiarchs; its sculpture changes with the body areas longitudinally from a smooth surface to a combination of longitudinal crests/ridges, grooves, irregular tubercles, and indistinct pits (Hemmings, 1978; Ivanov et al., 1996; Young, 1990).

In rhenanids, only the histology of *Ohioaspis* scales was investigated (Burrow & Turner, 1999). The middle layer is not developed as the main body of vascular canals is buried within the older generations of tubercles (Gross, 1973) and thus belongs to the superficial layer. The scales of *Ohioaspis* have stellate tubercles with a multitude of ridges around them (Burrow & Turner, 1999; Gross, 1973), whiles the scales of *Gemundina* and *Jagorina* are simply ornamented with round tubercles (Denison, 1978, figure 6C and D).

The absence of the spongiosa in the scale was ever perceived as a feature for acanthothoracids (Burrow, 1996; Denison, 1978). The middle layer is very weakly developed in scales of *Jerulalepis* and radotinids (Burrow, 1996, figure 3G; Stensiö, 1969, figure 255S–T). However, different conditions occur in the scales referred to *Murrindalaspis*, *Connemarraspis*, and *Romundina*, which bear a well-developed middle layer in addition to the complex ornamentation composed of stellate or polygonal tubercles as in the stem osteichthyan *Lophosteus* (Burrow, 2006; Burrow & Turner, 1998; Gross, 1969; Long & Young, 1988). Noteworthy is that the middle layer in the scales of *Romundina* exhibits two different conditions: poorly developed in some thin sections (Giles et al., 2013, figure 8A; Rucklin & Donoghue, 2015; Smith et al., 2017), and well-developed in other sections (Giles et al., 2013, figure 7F).

**Figure 8.**
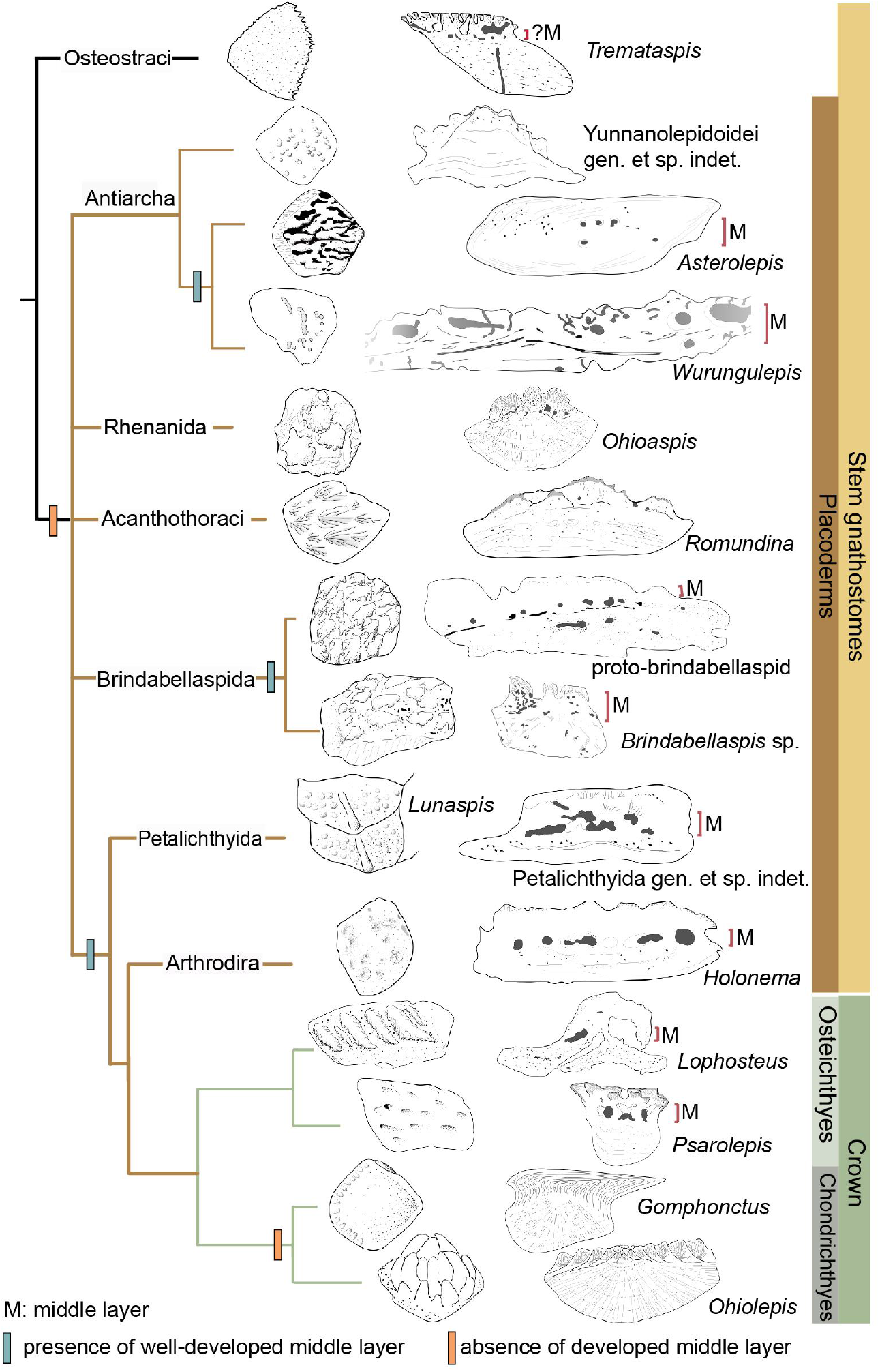
Sculpture pattern and histology of scales among gnathostomes. The phylogenetic hypothesis is after Zhu et al. (2016) as it accords with most of the recent hypotheses about the interrelationships of early jawed vertebrates. Taxon drawings are redrawn from source illustruations of Märss et al. (2015) for *Tremataspis*, Burrow and Turner (1999) and Ivanov et al. (1996) for *Asterolepis*, Burrow and Turner (1998) and Young (1990) for *Wurungulepis*, Giles et al. (2013) and Rucklin and Donoghue (2015) for *Romundina*, Burrow and Turner (1999) for *Ohioaspis*, Burrow and Turner (1999) for *Lunaspis*, Burrow and Turner (1998) for ‘proto-brindabellaspid’ and *Brindabellaspis*, Burrow et al. (2000) for Petalichthyida gen. et sp. indet., Trinajstic (1999) for *Holonema*, Gross (1969), Jerve et al. (2016) and Gross (1969) for *Lophosteus*, Qu et al. (2017) for *Psarolepis*, Sire et al. (2009) for *Gomphonctus*, and Gross (1973) for *Ohiolepis*.

In brindabellaspids, a type of scales with pavement/leaf-like tubercles was assigned to the ‘proto-brindabellaspid’ (Burrow & Turner, 1999). A thin layer of vascular canals, showing signs of centripetal apposition like osteons, represents the middle layer of the scale (Burrow & Turner, 1998). By comparison, the middle layer in the *Brindabellaspis* scale is much more developed (Burrow & Turner, 1999). Similarly, the ornamentation of the *Brindabellaspis* scale is more complicated than that of the ‘proto-brindabellaspid’ in bearing arrowhead-shaped tubercles (Burrow & Turner, 1998).

Petalichthyids and arthrodires constitute the successive sister groups of jawed stem gnathostomes, and their scales generally bear a well-developed middle layer (Burrow et al., 2000; Trinajstic, 1999), with a few exceptions of buchanosteid scales (Burrow & Turner, 1998). Concerning the ornamentation, the petalichthyid scales usually bear a diagonal ridge, surrounded by rounded or stellate tubercles (Burrow et al., 2000; Denison, 1978). The arthrodire scales are largely covered by rounded tubercles, or stellate tubercles that are either surrounded centrally by larger tubercles or combined with longitudinal ridges (Burrow & Turner, 1999; Johanson et al., 2019; Mark-Kurik & Young, 2003).

In crown gnathostomes, the middle layer occurs extensively in early osteichthyans, such as *Lophosteus* (Jerve et al., 2016) and *Psarolepis* (Qu et al., 2013; Schultze, 2016), but is widely absent in the chondrichthyan lineage including acanthodians (Andreev et al., 2016; Andreev et al., 2020; Burrow et al., 2020; Burrow et al., 2016; Chevrinais et al., 2017). The ornamentation on dermal scales seems slightly more variable and complex in osteichthyans than that of placoderms, as exampled by the *Andreolepis* and *Lophosteus* scales. *Andreolepis* usually carries parallel or united ridges along with grooves and pores (Chen et al., 2012), and *Lophosteus* carries serrated ridges (Jerve et al., 2016). The scale surface in the chondrichthyan lineage is often marked by pectinate ornamentation, consisting of raised ridges or denticulate ornament, occasionally accompanied by foramina.

The bipartite histological structure of the scale, comprising a thick basal layer and a superficial layer, was ever considered as a common feature for primitive placoderms (Burrow, 1996; Denison, 1978). New evidence from yunnanolepidoid scales supports this inference. Then the well-developed middle layer independently evolved several times in euantiarchs, brindabellaspids, some acanthothoracids, and a clade comprising more crownward placoderms including petalichthyids and arthrodires, plus crown gnathostomes, but lost again in the chondrichthyan total group. Another distinct trend is that the development of the well-developed middle layer is usually accompanied by the complication of ornamentation (*Figure 8*).

## Conclusions

A comprehensive study on the squamation of *P. xitunensis* using X-ray computed tomography and the histology of disarticulated scales provides the following new features in yunnanolepidoid antiarchs: 1) scales are generally rhombic, thick and show a concave base, but still vary in shape, size, ornamentation, concavity of the base, and overlap relationships in one individual, based on which we classified them into at least thirteen morphotypes for *P. xitunensis*; 2) nine areas of the post-thoracic body are distinguished to show the scale variations in the dorsal, flank, ventral, and caudal lobe regions, revealing the high regionalization of squamation at the root of jawed vertebrates; 3) yunnanolepidoid scales are histologically bipartite, composed of a thin upper superficial layer with multiple generations of tubercles and a thick basal layer.

Thin sections for yunnanolepidoid scales, together with evidence from other gnathostomes suggest that the well-developed middle cancellous layer of scales is primitively absent in jawed vertebrates, and independently evolved in euantiarchs, brindabellaspids, some acanthothoracids, petalichthyids, arthrodires, and crown gnathostomes. The development of the middle layer in the scales of early jawed vertebrates is accompanied by the complexity of the crown sculpture.

## Materials and methods

This study involves the holotype of *P. xitunensis* IVPP V11679.1 and six disarticulated scales IVPP V28642–V28647 (*Figure 6—figure supplement 6*) from Xitun village, Cuifengshan in Qujing city (Yunnan Province). They were collected from the middle part of the Xitun Formation (late Lochkovian) and housed in the Institute of Vertebrate Paleontology and Paleoanthropology.

V11679.1 was scanned using the Nano-CT system (Phoenix x-ray, GE Measurement & Control) at a pixel resolution of 5.99 μm in the Key Laboratory of Vertebrate Evolution and Human Origins of Chinese Academic of Science, Beijing. It was scanned at 90 kV, 100 μA, 1 s exposure time, with a 9 μm focal spot size. V28642–V28647 were processed with the diluted acetic acid and then scanned using the Nano-CT at 80 kV and100 μA, with a 1.83 μA voxel size. Raw data were pre-processed in VG Studio Max 3.3, and reconstructions were performed using Mimics 19.0.

### Institutional abbreviations

IVPP, Institute of Vertebrate Paleontology and Paleoanthropology, Chinese Academy of Sciences, Beijing, China.

### Anatomical abbreviations

ADL, anterior dorsolateral plate; adls, anterior dorsolateral scale; ads, anterior dorsal scale; afs, anterior flank scale; AMD, anterior median dorsal plate; AVL, anterior ventrolateral plate; avs: anterior ventral scute; b, base of the scale; bas, basal layer; c, crown of the scale; cls, caudal lobe scale; df, depressed field; dfs, dorsal fin scale; drs1–3, 1^st^, 2^nd^ and 3^rd^ dorsal ridge scales; fuls, posterodorsal fulcral scale; gr, groove on the ventral scute; gl, growth line; go1, go2, go3, first, second and third generations of odontodes; is, interspersed scale; lac, lacunae; lg, lateral line groove; ls.ad, anterodorsal lateral line scale; ls.ps, posterodorsal lateral line scale; PDL, posterior dorsolateral plate; pfs, posterior flank scale; PL, posterior lateral plate; PMD, posterior median dorsal plate; PVL, posterior ventrolateral plate; pvs, posterior ventral scute; rt, recrystallized tissue; sf, Sharpey’s fibers; sup, superficial layer; vls, ventrolateral scale.

### Anatomical abbreviations in supplementary figures

df, depressed field; dl, dorsal lamina of the anterior dorsolateral scale; dr, dorsal ridge on the third dorsal ridge scale; lc, lateral corner of the scale; lg, lateral line groove; p, pore; vl, ventral lamina of the anterior dorsolateral scale.

### Access to material

CT scan data (exported as *.mcs* files), reconstructions (exported as *.stl* files), and measurement data generated during this study will be available at https://doi.org/10.5061/dryad.fttdz08vb upon acceptance of the manuscript.

## Acknowledgements

We thank H. X. Lin (IVPP), Y. M. Hou (IVPP), and P. F. Yin (IVPP) for their assistance with CT scanning, and to L. T. Jia (IVPP) for his assistance with photographs. We also thank L. J. Peng (Qujing Normal University) for his help in making thin sections and Q. M. Qu (Xiamen University) for his constructive suggestions. This work was supported by the National Natural Science Foundation of China (42130209), and the Strategic Priority Research Program of Chinese Academy of Sciences (XDA19050102 and XDB26000000).

## Supplementary files

**Figure supplement 1.**
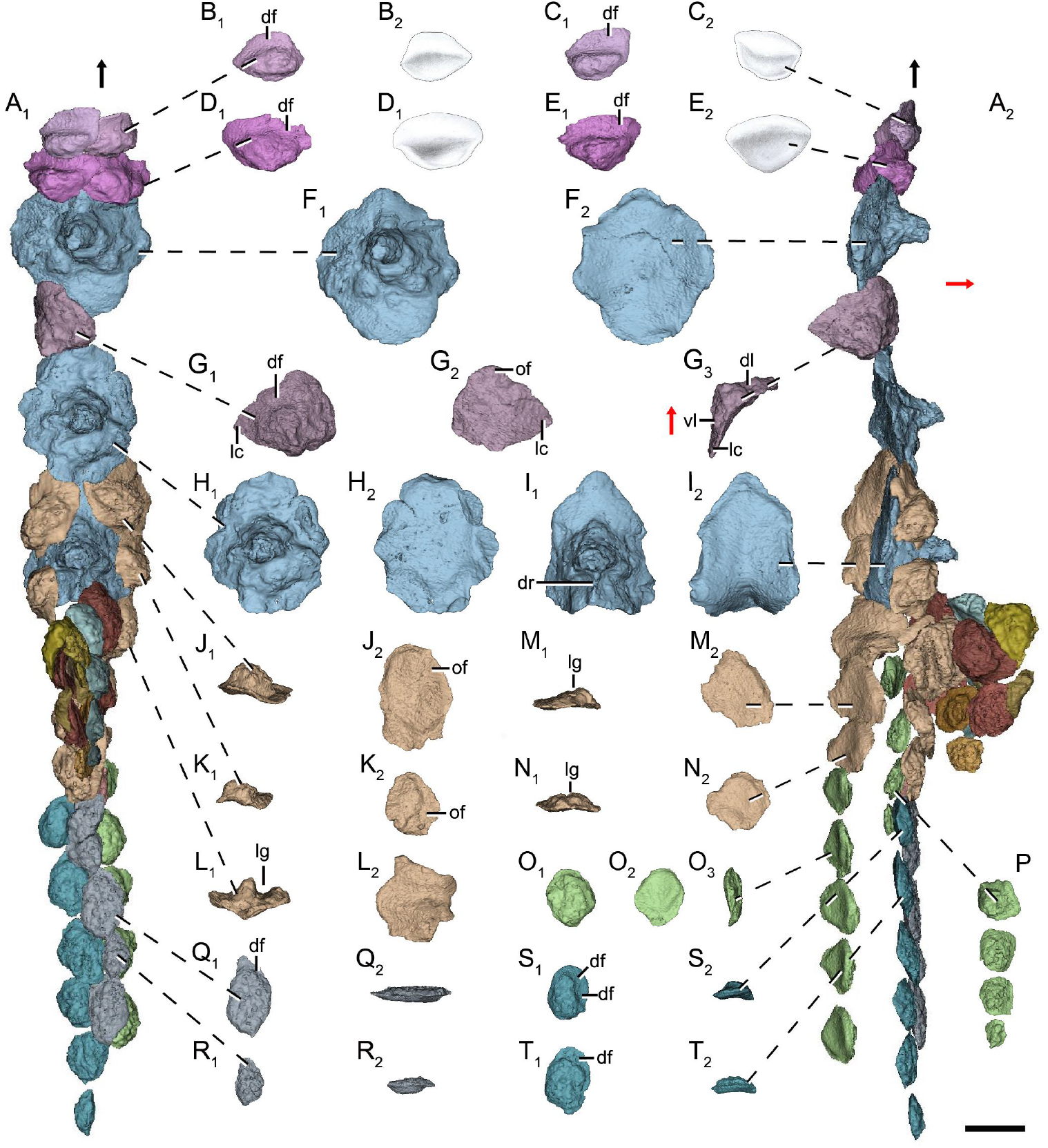
Dorsal scales of *Parayunnanolepis xitunensis* (IVPP V11679.1) based on CT scanning. (**A**) Dorsal squamation in (**A_1_**) dorsal and (**A_2_**) left lateral views. (**B**–**E**) Anterior dorsal scale in (**B_1_**–**E_1_**) dorsal and (**B**_2_–**E**_2_) ventral views. (**F**, **H** and **I**) First, second and third dorsal ridge scale in (**F_1_**, **H_1_** and **I_1_**) dorsal and (**F_2_**, **H_2_** and **I_2_**) ventral views. (**G**) Anterior dorsolateral scale in (**G_1_**) dorsal, (**G_2_**) ventral and (**G_3_**) posterior views. (**J**–**N**) anterodorsal lateral line scale in posterior (**J_1_**–**N_1_**) and ventral (**J_2_**–**N_2_**) views. (**O**) Posterodorsal lateral line scale in (**O_1_**) dorsal, (**O_2_**) ventral and (**O_3_**) left lateral views. (**P**) Posterodorsal lateral line scale preserved on the left side in dorsal view. (**Q**–**T**) Posterodorsal fulcral scale in (**Q_1_**–**T_1_**) dorsal, (**Q_2_**–**R_2_**) mesial and (**S_2_**–**T_2_**) posterior views. The black and red arrows indicate the anterior and dorsal directions, respectively. Scale bar equals to 1 mm.

**Figure supplement 2.**
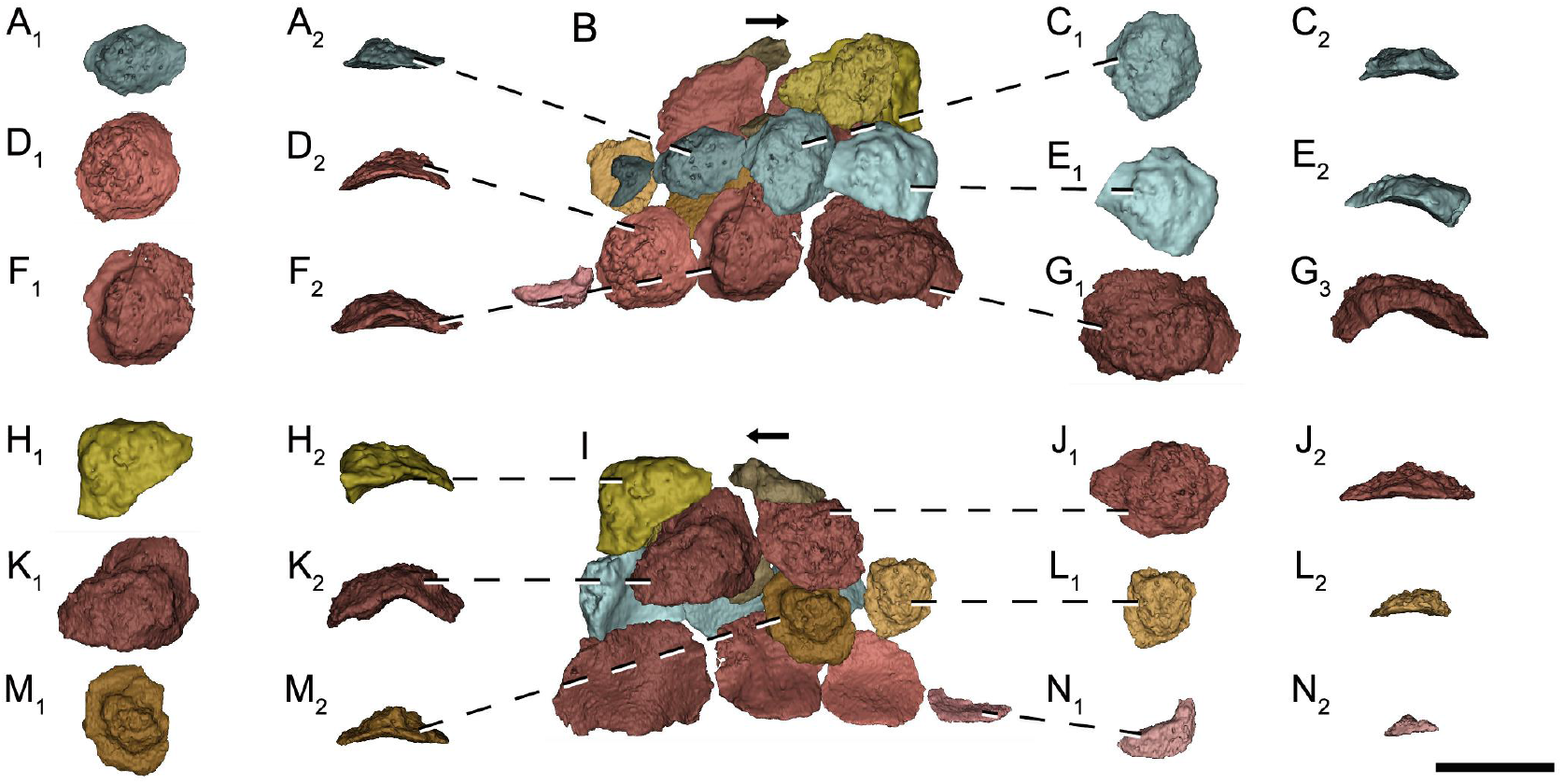
Dorsal fin scales of *Parayunnanolepis xitunensis* (IVPP V11679.1) based on CT scanning. (**A_1_, C_1_**–**H_1_, J_1_**–**N_1_**) Dorsal view. (**A_1_, C_1_**–**H_1_, J_1_**–**N_1_**) Ventral view. (**B** and **I**) Lateral view. The black arrow indicates the anterior direction. Scale bar equals to 1 mm.

**Figure supplement 3.**
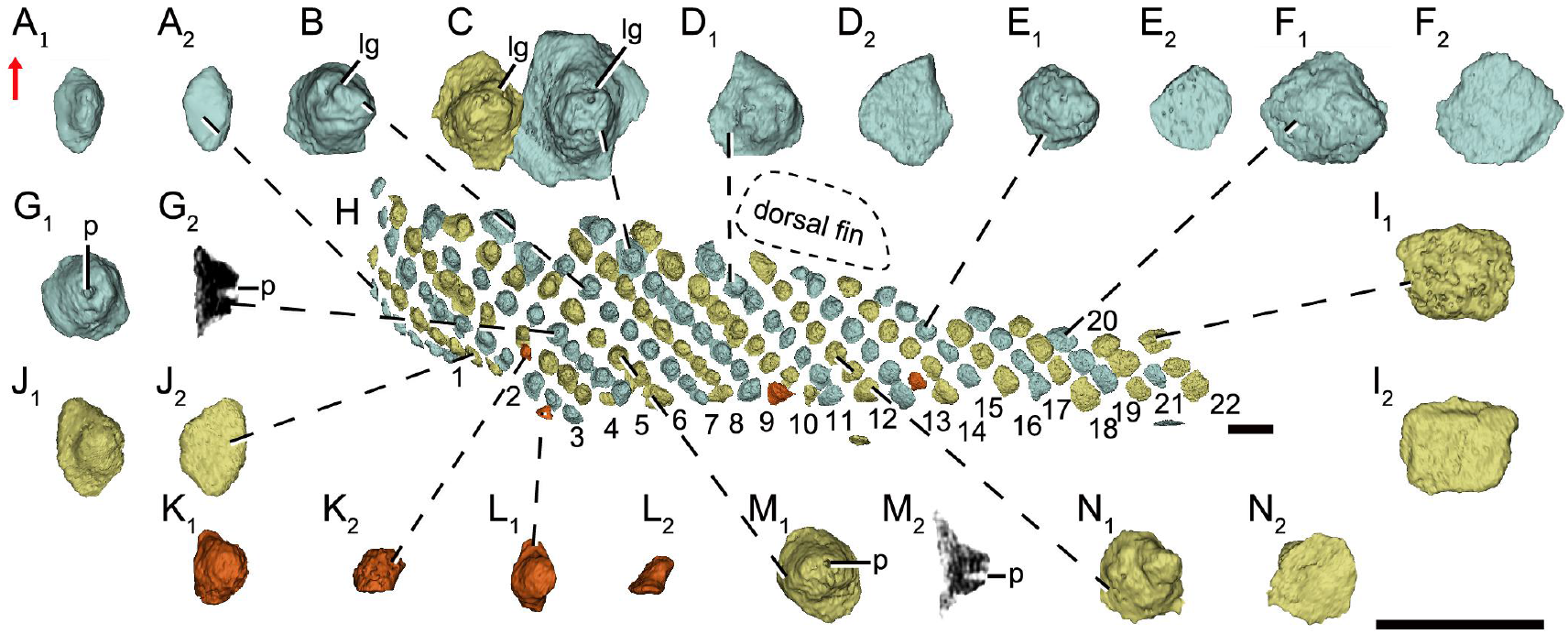
Flank scales on the left side of *Parayunnanolepis xitunensis* (IVPP V11679.1) based on CT scanning. (**A**–**E**, **G**, **J**–**N**) Anterior flank scales in dorsal (**A_1_**–**E_1_**, **G_1_**, **J_1_**–**N_1_**) and (**A_2_**–**E_2_**, **G_2_**, **J_2_**–**N_2_**) ventral views. (**E**, **F** and **I**) Posterior flank scales in (**E_1_**, **F_1_** and **I_1_**) dorsal and (**E_2_**, **F_2_** and **I_2_**) ventral views. (**H**) Flank squamation on the left side. The red arrow indicates the dorsal direction. Scale bars equal to 1 mm.

**Figure supplement 4.**
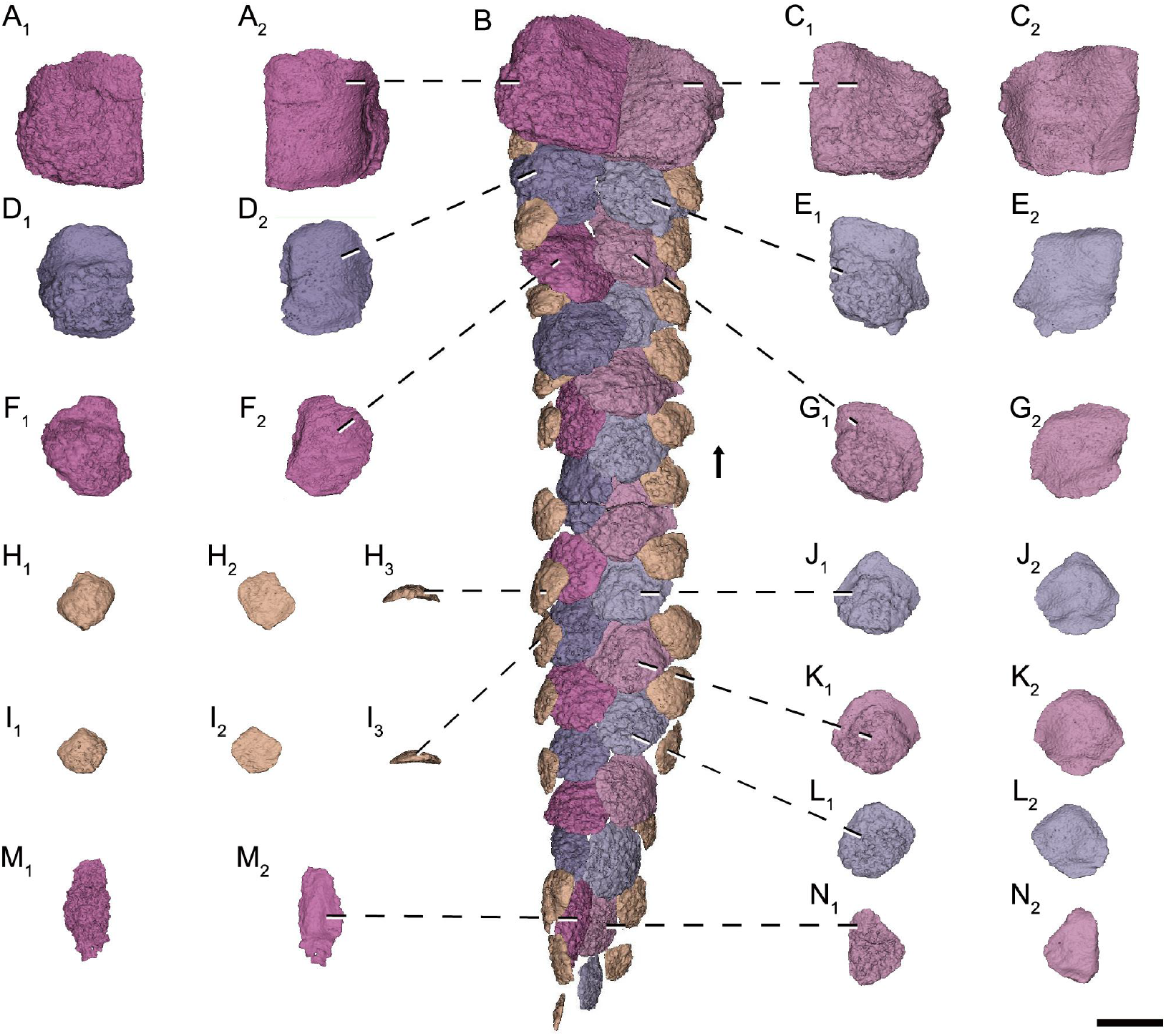
Ventral scales of *Parayunnanolepis xitunensis* (IVPP V11679.1) based on CT scanning. (**A**, and **C**–**G**) Anterior ventral scutes in (**A_1_**, and **C_1_**–**G_1_**) dorsal and (**A_2_**, and **C_2_**–**G_2_**) ventral views. (**B**) Ventral squaqmation in ventral view. (**H**–**I**) Ventrolateral scale in (**H_1_**–**I_1_**) dorsal, (**H_2_**–**I_2_**) ventral and (**H_3_**–**I_3_**) posterior views. (**J**–**N**) Posterior ventral scutes in (**J_1_**–**N_1_**) dorsal and (**J_2_**–**N_2_**) ventral views. The black arrow indicates the anterior direction. Scale bar equals to 1 mm.

**Figure supplement 5.**
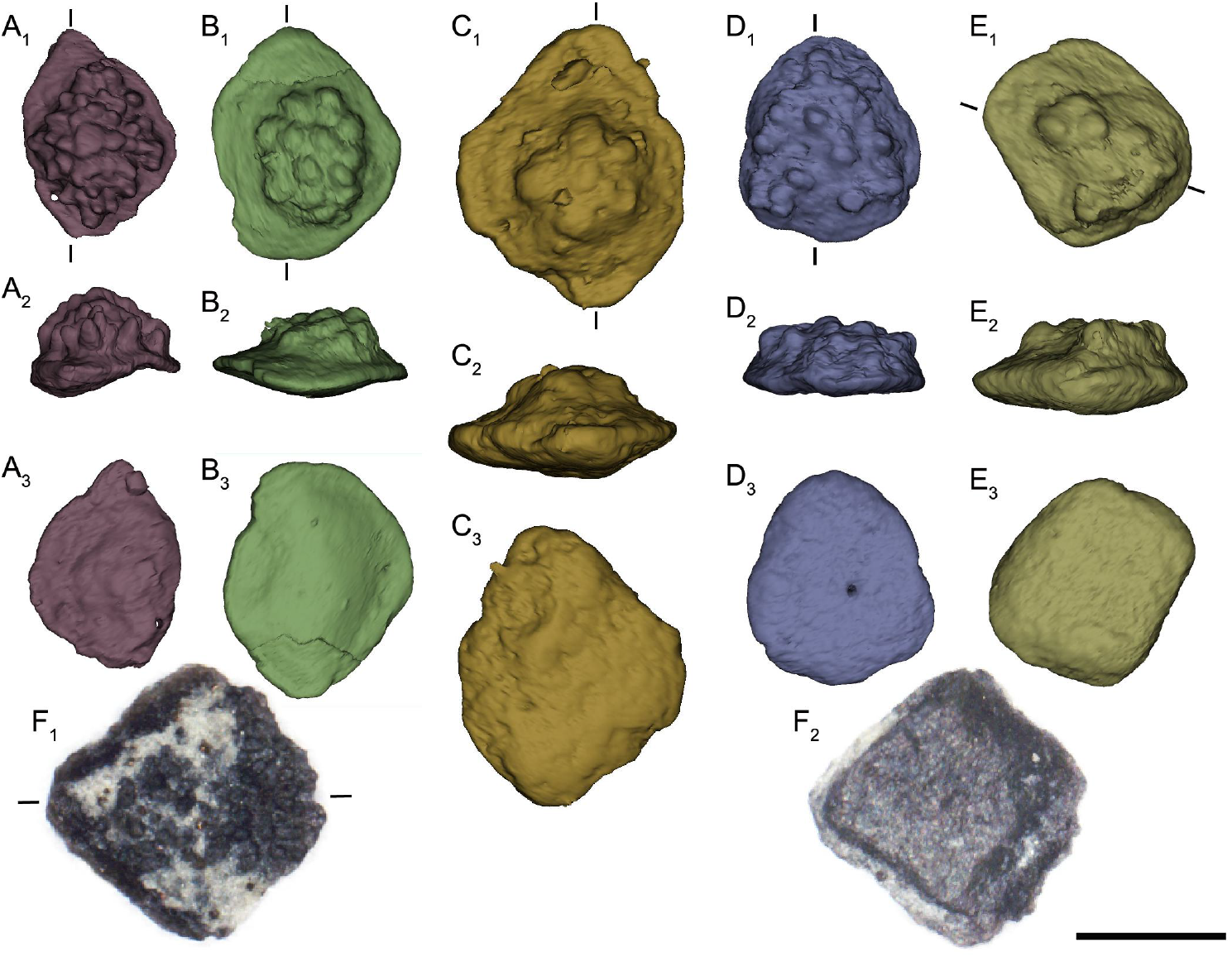
Yunnanolepidoid scales. (**A**–**E**) Reconstruction of IVPP V28642–V28647 based on CT scanning. (**A**) V28646. (**B**) V28643. (**C**) V28645. (**D**) V28644. (**E**) V28647. (**F**) V28642. Approximate planes of each histological section are indicated by short lines. Scale bar equals to 500 μm.

**Figure supplement 6.**
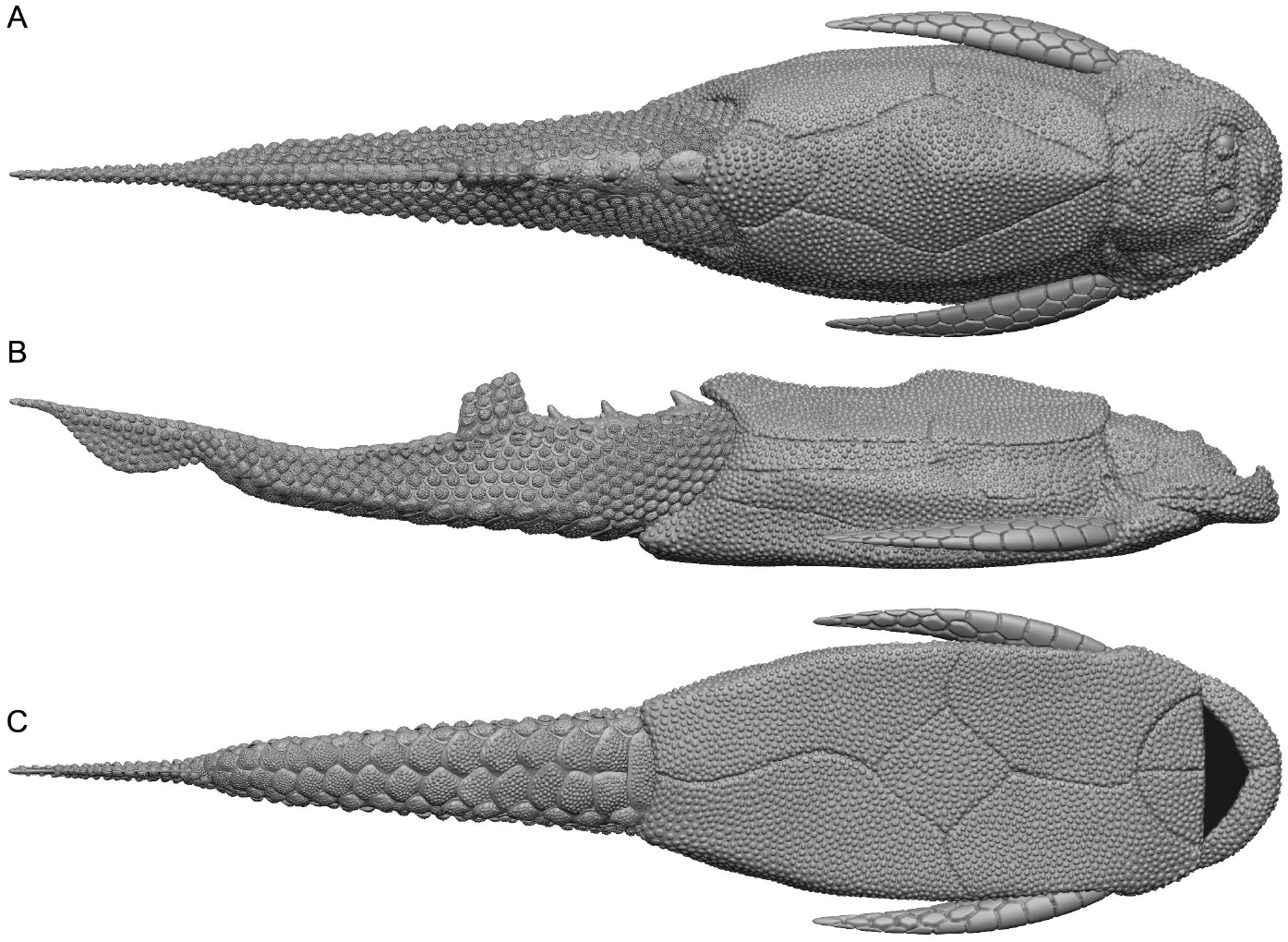
Reconstruction of *Parayunnanolepis xitunensis* by Dinghua Yang. (**A**) Dorsal view. (**B**) Lateral view. (**C**) Ventral view.

## Notes

### Competing Interest Statement

The authors have declared no competing interest.

## References

Andreev, P., Coates, M. I., Karatajute-Talimaa, V., Shelton, R. M., Cooper, P. R., Wang, N. Z., & Sansom, I. J. (2016). The systematics of the Mongolepidida (Chondrichthyes) and the Ordovician origins of the clade. PeerJ, 4, e1850. https://doi.org/10.7717/peerj.1850

Andreev, P. S., Zhao, W. J., Wang, N. Z., Smith, M. M., Li, Q., Cui, X. D., Zhu, M., & Sansom, I. J. (2020). Early Silurian chondrichthyans from the Tarim Basin (Xinjiang, China). PLoS One, 15(2), e0228589.

Brazeau, M. D., Giles, S., Dearden, R. P., Jerve, A., Ariunchimeg, Y., Zorig, E., Sansom, R., Guillerme, T., & Castiello, M. (2020). Endochondral bone in an Early Devonian ‘placoderm’ from Mongolia. Nature Ecology & Evolution, 4(11), 1477–1484. https://doi.org/10.1038/s41559-020-01290-2

Burrow, C. J., den Blaauwen, J., & Newman, M. (2020). A redescription of the three longest-known species of the acanthodian *Cheiracanthus* from the Middle Devonian of Scotland. Palaeontologia Electronica, 23(1), a15. https://doi.org/10.26879/1035

Burrow, C. J. (1996). Placoderm scales from the Lower Devonian of New South Wales, Australia. Modern Geology, 20, 351–369.

Burrow, C. J. (2006). Placoderm fauna from the Connemarra Formation (?late Lochkovian, Early Devonian), central New South Wales, Australia. Alcheringa, Special Issue 1, 59–88. https://doi.org/10.1080/03115510609506856

Burrow, C. J., den Blaauwen, J., Newman, M., & Davidson, R. (2016). The diplacanthid fishes (Acanthodii, Diplacanthiformes, Diplacanthidae) from the Middle Devonian of Scotland. Palaeontologia Electronica, 19(1), 1–83.

Burrow, C. J., & Turner, S. (1998). Devonian placoderm scales from Australia. Journal of Vertebrate Paleontology, 18(4), 677–695.

Burrow, C. J., & Turner, S. (1999). A review of placoderm scales, and their significance in placoderm phylogeny. Journal of Vertebrate Paleontology, 19(2), 204–219.

Burrow, C. J., Turner, S., & Wang, S. T. (2000). Devonian microvertebrates from Longmenshan, Sichuan, China: Taxonomic assessment. Courier Forschungsinstitut Senckenberg, 223, 391–451.

Chen, D. L., Janvier, P., Ahlberg, P. E., & Blom, H. (2012). Scale morphology and squamation of the Late Silurian osteichthyan *Andreolepis* from Gotland, Sweden. Historical Biology, 24(4), 411–423. https://doi.org/10.1080/08912963.2012.668187

Chevrinais, M., Sire, J. Y., & Cloutier, R. (2017). From body scale ontogeny to species ontogeny: Histological and morphological assessment of the Late Devonian acanthodian *Triazeugacanthus affinis* from Miguasha, Canada. PLoS One, 12(4), e0174655. https://doi.org/10.1371/journal.pone.0174655

Choo, B. (2012). Revision of the actinopterygian genus *Mimipiscis (=Mimia)* from the Upper Devonian Gogo Formation of Western Australia and the interrelationships of the early Actinopterygii. Earth and Environmental Science Transactions of the Royal Society of Edinburgh, 102(02), 77–104. https://doi.org/10.1017/s1755691011011029

Choo, B., Long, J. A., & Trinajstic, K. (2009). A new genus and species of basal actinopterygian fish from the Upper Devonian Gogo Formation of Western Australia. Acta Zoologica, 90, 194–210. https://doi.org/10.1111/j.1463-6395.2008.00370.x

Choo, B., Zhu, M., Qu, Q. M., Yu, X. B., Jia, L. T., & Zhao, W. J. (2017). A new osteichthyan from the late Silurian of Yunnan, China. PLoS One, 12(3), e0170929. https://doi.org/10.1371/journal.pone.0170929

Cui, X. D., Qiao, T., & Zhu, M. (2019). Scale morphology and squamation pattern of *Guiyu oneiros* provide new insights into early osteichthyan body plan. Scientific Reports, 9(1), 4411. https://doi.org/10.1038/s41598-019-40845-7

Cui, X. D., Qu, Q. M., Andreev, P. S., Li, Q., Mai, H. J., & Zhu, M. (2021). Modeling scale morphogenesis in a Devonian chondrichthyan and scale growth patterns in crown gnathostomes. Journal of Vertebrate Paleontology, 41(2), e1930018. https://doi.org/10.1080/02724634.2021.1930018

Dearden, R. P., Stockey, C., & Brazeau, M. D. (2019). The pharynx of the stem-chondrichthyan *Ptomacanthus* and the early evolution of the gnathostome gill skeleton. Nature Communications, 10(1), 2050. https://doi.org/10.1038/s41467-019-10032-3

Denison, R. (1978). Placodermi. Gustav Fischer Verlag.

Donoghue, P. C. J., & Sansom, I. J. (2002). Origin and early evolution of vertebrate skeletonization. Microscopy Research and Technique, 59(5), 352–372. https://doi.org/10.1002/jemt.10217

Downs, J. P., & Donoghue, P. C. J. (2009). Skeletal histology of *Bothriolepis canadensis* (Placodermi, Antiarchi) and evolution of the skeleton at the origin of jawed vertebrates. Journal of Morphology, 270(11), 1364–1380. https://doi.org/10.1002/jmor.10765

Dupret, V., Sanchez, S., Goujet, D., Tafforeau, P., & Ahlberg, P. E. (2014). A primitive placoderm sheds light on the origin of the jawed vertebrate face. Nature, 507, 500–503. https://doi.org/10.1038/nature12980

Ferron, H. G., & Botella, H. (2017). Squamation and ecology of thelodonts. PLoS One, 12(2), e0172781. https://doi.org/10.1371/journal.pone.0172781

Giles, S., Friedman, M., & Brazeau, M. D. (2015). Osteichthyan-like cranial conditions in an Early Devonian stem gnathostome. Nature, 520, 82–85. https://doi.org/10.1038/nature14065

Giles, S., Rücklin, M., & Donoghue, P. C. J. (2013). Histology of “placoderm” dermal skeletons: Implications for the nature of the ancestral gnathostome. Journal of Morphology, 274(6), 627–644. https://doi.org/10.1002/jmor.20119

Goujet, D. F. (1973). *Sigaspis*, un nouvel arthrodire du Dévonien inférieur du Spitsberg. Palaeontographica Abteilung A, 143, 73–88.

Gross, W. (1963). *Gemuendina stuertzi* Traquair. Neuuntersuchung. Notizblatt des hessisches Landesanstalt für Bodenforschung, 91, 36–73.

Gross, W. (1969). *Lophosteus superbus* Pander, ein Teleostome aus dem Silur Oesels. Lethaia, 2, 15–47.

Gross, W. (1973). Kleinschuppen, Flossenstacheln und Zähne von Fischen aus europäischen und nordamerikanischen Boncbeds des Devons. Palaeontographica Abteilung A, 142, 51–155.

Hemmings, S. K. (1978). The Old Red Sandstone antiarchs of Scotland: *Pterichthyodes* and *Microbrachius*. Monographs of the Palaeontographical Society, 131, 1–64.

Ivanov, A., Cherepanov, G., & Luksevics, E. (1995). Ontogenetic development of antiarch dermal ossifications. Geobios Mémoire Spécial, 19, 97–102.

Ivanov, A., Lukševics, E., & Upeniece, I. (1996). The squamous part of an asterolepid body. Modern Geology, 20, 399–410.

Janvier, P. (1996). Early Vertebrates. Clarendon Press.

Jerve, A., Qu, Q. M., Sanchez, S., Ahlberg, P. E., & Haitina, T. (2017). Vascularization and odontode structure of a dorsal ridge spine of *Romundina stellina* Ørvig 1975. PLoS One, 12(12), e0189833.

Jerve, A., Qu, Q. M., Sanchez, S., Blom, H., & Ahlberg, P. E. (2016). Three-dimensional paleohistology of the scale and median fin spine of *Lophosteus superbus* (Pander 1856). PeerJ, 4, e2521. https://doi.org/10.7717/peerj.2521

Johanson, Z. (1997). New *Remigolepis* (Placodermi; Antiarchi) from Canowindra, New South Wales, Australia. Geological Magazine, 134(6), 813–846.

Johanson, Z., Underwood, C., & Richter, M. (Eds.). (2019). Evolution and Development of Fishes. Cambridge University Press. https://doi.org/10.1017/9781316832172.

Li, Q., Zhu, Y. A., Lu, J., Chen, Y., Wang, J. H., Peng, L. J., Wei, G. B., & Zhu, M. (2021). A new Silurian fish close to the common ancestor of modern gnathostomes. Current Biology. https://doi.org/10.1016/j.cub.2021.05.053

Long, J. A. (1983). New bothriolepid fish from the late Devonian of Victoria, Australia. Palaeontology, 26(2), 295–320.

Long, J. A., Mark-Kurik, E., Johanson, Z., Lee, M. S., Young, G. C., Zhu, M., Ahlberg, P. E., Newman, M., Jones, R., Blaauwen, J. D., Choo, B., & Trinajstic, K. (2015). Copulation in antiarch placoderms and the origin of gnathostome internal fertilization. Nature, 517, 196–199. https://doi.org/10.1038/nature13825

Long, J. A., & Werdelin, L. (1986). A new Late Devonian bothriolepid (Placodermi, Antiarcha) from Victoria, with descriptions of other species from the state. Alcheringa, 10(4), 355–399.

Long, J. A., & Young, G. C. (1988). Acanthothoracid remains from the Early Devonian of New South Wales, including a complete sclerotic capsule and pelvic girdle. Memoirs of the Association of Australasian Palaeontologists, 7, 65–80.

Lukševičs, E. (1991). New *Remigolepis* (Pisces, Antiarchi) from the Famennian deposits of the Central Devonian Field (Russia, Tula region). Daba un muzejs, 3, 51–56.

Lyarskaya, L. (1977). Noviye danniye o *Asterolepis ornata* Eichwald iz rannefranskih otlozheniy Pribaltiki. In M. V.V. (Ed.), Notes on phylogeny and systematic of fossil fish and agnathans (pp. 36–44). Nauka publ.

Mark-Kurik, E., & Young, G. C. (2003). A new buchanosteid arthrodire (placoderm fish) from the Early Devonian of the Ural Mountains. Journal of Vertebrate Paleontology, 23(1), 13–27.

Märss, T., Afanassieva, O., & Blom, H. (2015). Biodiversity of the Silurian osteostracans of the East Baltic. Earth and Environmental Science Transactions of the Royal Society of Edinburgh, 105(02), 73–148. https://doi.org/10.1017/s1755691014000218

Ørvig, T. (1968). The dermal skeleton, general considerations. In T. Ørvig (Ed.), Current Problems of Lower Vertebrate Phylogeney. Nobel Symposium 4 (pp. 373–398). Almqvist & Wiksell.

Ørvig, T. (1975). Description, with special reference to the dermal skeleton, of a new radotinid arthrodire from the Gedinnian of Arctic Canada. In J. P. Lehman (Ed.), Problèmes actuels de Paléontologie-Evolution des Vertébrés (Vol. 218, pp. 41–71). Colloques Internationaux du Centre National de la Recherche Scientifique.

Pan, J., Huo, F. C., Cao, J. X., Gu, Q. C., Liu, S. Y., Wang, J. Q., Gao, L. D., & Liu, C. (1987). Continental Devonian System of Ningxia and its Biotas. Geological Publishing House.

Qiao, T., King, B., Long, J. A., Ahlberg, P. E., & Zhu, M. (2016). Early gnathostome phylogeny revisited: multiple method consensus. PLoS One, 11(9), e0163157. https://doi.org/10.1371/journal.pone.0163157

Qu, Q. M., Blom, H., Sanchez, S., & Ahlberg, P. (2015). Three-dimensional virtual histology of Silurian osteostracan scales revealed by synchrotron radiation microtomography. Journal of Morphology, 276(8), 873–888. https://doi.org/10.1002/jmor.20386

Qu, Q. M., Haitina, T., Zhu, M., & Ahlberg, P. E. (2015). New genomic and fossil data illuminate the origin of enamel. Nature, 526(7571), 108–111. https://doi.org/10.1038/nature15259

Qu, Q. M., Sanchez, S., Zhu, M., Blom, H., & Ahlberg, P. E. (2017). The origin of novel features by changes in developmental mechanisms: ontogeny and three-dimensional microanatomy of polyodontode scales of two early osteichthyans. Biological Reviews, 92(2), 1189–1212. https://doi.org/10.1111/brv.12277

Qu, Q. M., Zhu, M., & Wang, W. (2013). Scales and dermal skeletal histology of an early bony fish *Psarolepis romeri* and their bearing on the evolution of rhombic scales and hard tissues. PLoS One, 8(4), e61485. https://doi.org/10.1371/journal.pone.0061485.g001

Reif, W. E. (1985). Squamation and ecology of sharks. Courier Forschungsinstitut Senckenberg, 78, 1–255.

Ribeiro, A. C., Ribeiro, G. C., Varejão, F. G., Battirola, L. D., Pessoa, E. M., Simões, M. G., Warren, L. V., Riccomini, C., & Poyato-Ariza, F. J. (2021). Towards an actualistic view of the Crato Konservat-Lagerstätte paleoenvironment: A new hypothesis as an Early Cretaceous (Aptian) equatorial and semi-arid wetland. Earth-Science Reviews, 103573. http://doi.org/10.1016/j.earscirev.2021.103573

Rucklin, M., & Donoghue, P. C. (2015). *Romundina* and the evolutionary origin of teeth. Biology Letters, 11(6), 20150326. https://doi.org/10.1098/rsbl.2015.0326

Schultze, H. P. (2016). Scales, enamel, cosmine, ganoine, and early osteichthyans. Comptes Rendus Palevol, 15(1–2), 83–102. https://doi.org/10.1016/j.crpv.2015.04.001

Sire, J. Y., Donoghue, P. C. J., & Vickaryous, M. K. (2009). Origin and evolution of the integumentary skeleton in non-tetrapod vertebrates. Journal of Anatomy, 214(4), 409–440. https://doi.org/10.1111/j.1469-7580.2009.01046.x

Smith, M. M., Clark, B., Goujet, D., Johanson, Z., & Smith, A. (2017). Evolutionary origins of teeth in jawed vertebrates: conflicting data from acanthothoracid dental plates (‘Placodermi’). Palaeontology, 60(6), 829–836. https://doi.org/10.1111/pala.12318

Stensiö, E. (1969). Elasmobranchiomorphi Placodermata Arthrodires. In J. Piveteau (Ed.), Traité de Paléontologie (Vol. 4(2), pp. 71–692). Masson.

Stensiö, E. A. (1948). On the Placodermi of the Upper Devonian of East Greenland. II. Antiarchi: subfamily Bothriolepinae. With an attempt at a revision of the previously described species of that family. Meddelelser om Grønland, 139, 1–622.

Trinajstic, K. (1999). New anatomical information on Holonema (Placodermi) based on material from the Frasnian Gogo Formation and the Givetian-Frasnian Gneudna Formation, Western Australia. New information on placoderms from Late Devonian of Western Australia, 21(1), 69–84.

Turner, S. (1991). Monophyly and interrelationships of the Thelodonti. In M. M. Chang, Y. H. Liu, & G. R. Zhang (Eds.), Early Vertebrates and Related Problems of Evolutionary Biology (pp. 87–119). Science Press.

Upeniece, I., & Upenieks, J. (1992). Young Upper Devonian antiarch *(Asterolepis)* individuals from the Lode quarry, Latvia. In E. Mark-Kurik (Ed.), Fossil Fishes as Living Animals (pp. 167–176). Academy of Sciences of Estonia.

Vaškaninová, V., Chen, D. L., Tafforeau, P., Johanson, Z., Ekrt, B., Blom, H., & Ahlberg, P. E. (2020). Marginal dentition and multiple dermal jawbones as the ancestral condition of jawed vertebrates. Science, 369, 211–216. https://doi.org/10.1126/science.aaz9431

Young, G. C. (1990). New antiarchs (Devonian placoderm fishes) from Queensland, with comments on placoderm phylogeny and biogeography. Memoirs of the Queensland Museum, 28(1), 35–50.

Young, G. C. (2003). Did placoderm fish have teeth? Journal of Vertebrate Paleontology, 23(4), 987–990.

Zhang, G. R. (1978). The antiarchs from the Early Devonian of Yunnan. Vertebrata PalAsiatica, 16(3), 147–186.

Zhang, G. R., Wang, J. Q., & Wang, N. Z. (2001). The structure of pectoral fin and tail of Yunnanolepidoidei, with a discussion of the pectoral fin of chuchinolepids. Vertebrata PalAsiatica, 39(1), 1–13.

Zhao, W. J., & Zhu, M. (2010). Siluro-Devonian vertebrate biostratigraphy and biogeography of China. Palaeoworld, 19(1–2), 4–26. https://doi.org/10.1016/j.palwor.2009.11.007

Zhu, M. (1996). The phylogeny of the Antiarcha (Placodermi, Pisces), with the description of Early Devonian antiarchs from Qujing, Yunnan, China. Bulletin du Muséum nationa1 d’Histoire naturelle, 18, 233–347.

Zhu, M., Ahlberg, P. E., Pan, Z. H., Zhu, Y. A., Qiao, T., Zhao, W. J., Jia, L. T., & Lu, J. (2016). A Silurian maxillate placoderm illuminates jaw evolution. Science, 354(6310), 334–336. https://doi.org/10.1126/science.aah3764

Zhu, M., Wang, N. Z., & Wang, J. Q. (2000). Devonian macro- and microvetebrate assemblages of China. Courier Forschungsinstitut Senckenberg, 223, 361–372.

Zhu, M., Yu, X., Ahlberg, P. E., Choo, B., Lu, J., Qiao, T., Qu, Q. M., Zhao, W. J., Jia, L. T., Blom, H., & Zhu, Y. A. (2013). A Silurian placoderm with osteichthyan-like marginal jaw bones. Nature, 502(7470), 188–193. https://doi.org/10.1038/nature12617

Zhu, M., Yu, X., Choo, B., Wang, J. Q., & Jia, L. T. (2012). An antiarch placoderm shows that pelvic girdles arose at the root of jawed vertebrates. Biology Letters, 8(3), 453–456. https://doi.org/10.1098/rsbl.2011.1033

Zhu, M., Zhao, W. J., Jia, L. T., Lu, J., Qiao, T., & Qu, Q. M. (2009). The oldest articulated osteichthyan reveals mosaic gnathostome characters. Nature, 458(7237), 469–474. https://doi.org/ https://doi.org/10.1038/nature07855

Zhu, Y. A., Giles, S., Young, G. C., Hu, Y., Bazzi, M., Ahlberg, P. E., Zhu, M., & Lu, J. (2021). Endocast and bony labyrinth of a Devonian “placoderm” challenges stem gnathostome phylogeny. Current Biology. https://doi.org/10.1016/j.cub.2020.12.046

